# Manifolds for the neural representation of 3D spatial experience

**DOI:** 10.64898/2026.10.22.760836

**Authors:** Raymond Doudlah, Mengyu Xiao, Satchal Postlewaite, Byounghoon Kim, Noah J. Cowan, Ari Rosenberg

**Affiliations:** Department of Neuroscience, School of Medicine and Public Health, University of Wisconsin–Madison, Madison, WI 53705, USA; Department of Mechanical Engineering, Johns Hopkins University, Baltimore, MD 21218, USA; Department of Neurobiology, University of Chicago, Chicago, IL 60637, USA

**Keywords:** neural manifolds, topology, 3D vision, navigation, head direction

## Abstract

The precision with which animals navigate and interact with objects depends in part on how their brains represent position and orientation in three dimensions. A central challenge is to explain within a common framework how neural systems ranging from insects to primates represent three-dimensional (3D) pose. Here we develop a framework based on the hypothesis that neural representations of pose lie on manifolds derived from the Special Euclidean Group, SE(3), which describes the full space of rigid-body positions and orientations in three dimensions. The framework shows how movement constraints, object geometry, and visual features jointly determine the topology of the resulting pose manifold, yielding a taxonomy that guides experimental tests. We apply the framework to derive low-dimensional manifolds for neural representations of 3D object orientation in macaque parietal cortex and head direction in the limbic systems of fruit bats, mice, and monkeys. This analysis also guides the construction of neuronal tuning functions that respect the topology determined by experimental conditions. Together, these results provide a common approach for linking the geometry of 3D pose to neural coding across species and brain systems.

**Significance statement:** Behaviors such as reaching and grasping are guided by visual estimates of 3D object pose, whereas postural control and navigation depend on multisensory estimates of how the self is posed in the world. Despite being resolved by distinct neural circuits, we propose that these computational goals share a common topology which we use to develop a unified mathematical framework for describing 3D spatial experience. The framework accounts for neuronal representations found across distinct sensorimotor and cognitive systems, and multiple species. Through the application of geometric topology, this work reveals formal equivalencies across disparate neural systems and research domains, and outlines a deductive strategy for testing theories of neuronal coding.

**Declaration of competing interests:** The authors declare no competing interests.

## Introduction

Complex behaviors such as tool use and navigation require representations of the position and orientation of objects and self, suggesting that neural systems may have evolved representations on manifolds that reflect the underlying geometry and topology of 3D pose. Indeed, seminal discoveries in systems neuroscience included mechanisms of pose selectivity. For instance, primary visual cortex (V1) encodes the two-dimensional (2D) pose of local edges (Hubel and Wiesel 1959), motor cortex encodes planar reach direction (Georgopoulos et al., 1982), and limbic structures contain neurons selective for head direction in the horizontal plane (Taube et al., 1990). These studies of 2D selectivity laid the foundation for investigating how neurons encode the full six degrees of freedom (DOFs) that specify position and orientation in 3D space (Craig, 2005). Determining how those six DOFs are encoded drives visual (Orban, 2011; Rosenberg et al., 2023, 2025), auditory (Knudsen and Konishi, 1978; Carlini et al., 2024), haptic (Lutz and Bensmaia, 2021), motor and postural control (Murata et al., 2000; Galletti et al., 2003; Theys et al., 2012), and spatial navigation (Finkelstein et al., 2015; Angelaki et al., 2020; Williams et al., 2026) research.

Neuronal coding theories have long been grounded in the analysis of manifolds – often used to describe lower-dimensional structure within higher-dimensional spaces. Early approaches derived manifolds describing neuronal feature selectivity and then related the geometries to the cortical organization of those features (Swindale, 1996). For example, the orientation and phase selectivity of V1 neurons is described by the geometry of a Klein bottle (Tanaka, 1995), accounting for the pinwheel organization of orientation maps (Bonhoeffer and Grinvald, 1991). The representation of orientation and spatial frequency in V1 on a spherical manifold (Bressloff and Cowan, 2003) likewise predicts the co-localization of orientation pinwheels and extreme spatial frequencies (Issa et al., 2000), as well as broader cortical activity patterns (Singh et al., 2008).

High-density electrodes and cellular imaging have since shifted the field toward data-driven inductive approaches that use dimensionality reduction to infer manifolds from neural recordings (Cunningham and Yu, 2014; Seelig and Jayaraman, 2015; Langdon et al., 2023; Curto and Sanderson, 2025) (Fig. 1A). The approach has facilitated discovery across multiple domains (DiCarlo et al., 2012; Gallego et al., 2017); however, the need for large, unbiased datasets makes it costly and time consuming. It is also inherently *post hoc*, potentially limiting the generality of particular findings.

**Figure 1.**
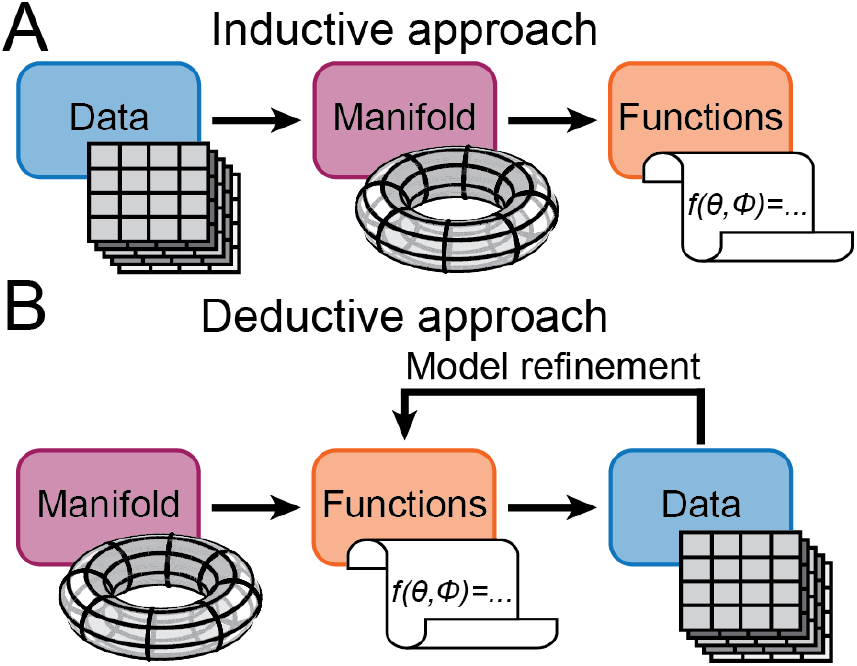
Inductive vs. deductive strategies. **(A)** Inductive strategy. First, substantial data collection occurs. Second, data reduction techniques are used to infer an underlying manifold. Third, tuning functions are determined and neuronal codes deciphered. **(B)** Deductive strategy. First, a manifold structure is derived from first principles. Second, tuning functions operating on the manifold are determined and fit to data, empowering model refinement. Third, neuronal codes are deciphered.

Here we introduce a deductive approach grounded in the topology of 3D rigid-body transformations, the Special Euclidean Group, SE(3). After introducing the full six-dimensional pose space, we use examples to show how movement constraints, object geometry, and visual features jointly determine the topology of pose manifolds and what this implies for neuronal coding. We derive tuning functions that respect these topologies and use them to show that object orientation and head direction selectivity in cortical and subcortical areas of bats, mice, and monkeys match the theoretical predictions. Across domains, we show that deductive and inductive approaches converge on equivalent manifolds. This work provides a unified approach for deriving falsifiable predictions about neural representations of 3D pose and making the consequences of experimental design explicit.

## Results

This section provides a broadly accessible introduction to key mathematical concepts and their connections to neuronal coding.

A manifold describes the latent geometric structure of a set of variables, such as distances, angles, or neuronal firing rates. A configuration space is the set of all possible states of a system, such as the positions and orientations of an object. Many configuration spaces are also manifolds: spaces that locally resemble a line, a plane, or a higher-dimensional Euclidean space, but can have more complex global structures. Lines and circles are both 1D manifolds, but whereas a line is globally a line (ℝ^1^), a circle requires at least two dimensions for a Euclidean embedding (2D Euclidean space, ℝ^2^; Supplementary Information Fig. **??**A). Similarly, a sphere is a 2D manifold embedded in 3D Euclidean space (ℝ^3^; Supplementary Information Fig. **??**B). Thus, a manifold’s dimensionality is that of the Euclidean space describing its local structure, not the space in which it is embedded. Relevant manifolds in this paper and their geometries are summarized in Table 1.

**Table 1.** Manifolds and geometric interpretations.

| Notation | Manifold dimension | Representation coordinates | Geometric interpretation |
| --- | --- | --- | --- |
| $\mathbb{R}^1$ | 1D | 1D | 1D Euclidean space; Line |
| $\mathbb{R}^2$ | 2D | 2D | 2D Euclidean space; Plane |
| $\mathbb{R}^3$ | 3D | 3D | 3D Euclidean space |
| $S^1$ | 1D | 2D | Unit circle |
| $S^2$ | 2D | 3D | Unit sphere |
| $S^3$ | 3D | 4D | Unit 3-sphere |
| $T^2$ | 2D | 3D | Torus ($S^1 \times S^1$) |
| $SO(3)$ | 3D | 4D | $S^3/(\mathbf{x} \sim -\mathbf{x})$ |
| $SE(3)$ | 6D | 7D | $\mathbb{R}^3 \times SO(3)$ |

Manifolds are foundational to neural coding because their geometries constrain the properties of functions (Hirsch, 1976; Mardia and Jupp, 2000). Consider an object’s positive distance *d* from an observer and its direction of planar motion. Excluding coincidence of the observer and object, *d* ∈ (0, ∞). Although distance is a positive real coordinate, (0, ∞) is diffeomorphic to ℝ^1^, for example under the map *d* ↦ log *d*. Thus, distance lies on the 1D manifold ℝ^1^, whereas direction of planar motion lies on the circular manifold S^1^. These geometries imply that, in a neural population encoding these quantities, distance tuning curves are functions over ℝ^1^, whereas direction tuning curves are functions over S^1^. The joint configuration space is the Cartesian product ℝ^1^ × S^1^ and is geometrically an open-ended cylinder (locally planar; Supplementary Information Fig. **??**C). A slice through the cylinder, S^1^, describes all directions at a fixed distance and a lengthwise cut, ℝ^1^, describes all distances for a fixed direction of motion. In turn, the cylinder is a submanifold of a higher-dimensional manifold representing all features encoded by the neuronal population (position, direction, speed, etc.).

### The geometry of 3D spatial experience

Representing the full dimensionality of spatial experience requires neural systems to encode the three translational DOFs specifying position and three rotational DOFs specifying orientation. These 6 DOFs collectively define 3D pose (Craig, 2005).

Position is parameterized by a 3-vector, **p** ∈ ℝ^3^, where each element is a location along one dimension. Orientation is described by a rotation matrix, *R*, that preserves vector length and the handedness of the reference frame. Length preservation requires the columns of *R* to form an orthonormal basis, *R*^*⊤*^*R* = *I* (identity matrix). Preservation of handedness requires det *R* = +1. In ℝ^3^, the group of 3 *×* 3 matrices satisfying these constraints is the Special Orthogonal Group, SO(3) = {*R* ∈ ℝ^3*×*3^ : *R*^*⊤*^*R* = *I*, det *R* = +1}.

Interpreting how translations and rotations change an object’s pose requires the definition of a coordinate frame, rotation sequence, and reference orientation. Because different conventions are formally equivalent, they are often selected for computational convenience or historical reasons. Two conventions used in research explored here are illustrated in Fig. 2A,B.

**Figure 2.**
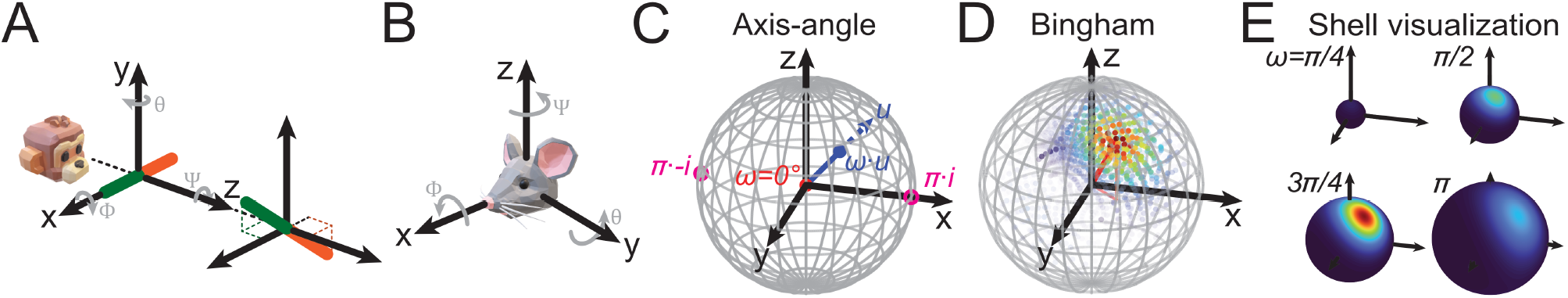
Coordinate frames and visualization of functions on SO(3). **(A)** Left-handed frame describing the (*x, y, z*) position and (*ϕ, θ, ψ*) orientation of a viewed object. *Left:* A bar in its reference orientation. *Right:* Orientation after applying *ϕ* = 20° and *ψ* = 225°. **(B)** Right-handed frame describing head direction. **(C)** Axis-angle visualization of SO(3) = S^3^*/*(**q** *∼ ™***q**). Data are projected onto a solid sphere of radius *π* in R^3^. Each orientation is represented by a point *ω · u*, where *ω* is the rotation angle (0 *≤ ω ≤ π*), and the unit vector *u* ∈ R^3^ is the rotation axis (blue). The reference orientation (*ω* = 0°) is at the center of the sphere (red). The outer shell is antipodally symmetric such that +*πu* and *™πu* are the same orientation (pink). **(D)** Example Bingham function. Firing rates are indicated by color and opacity to “see” into the sphere and minimize occlusion since it is a solid object. Higher (lower) rates are red (blue) and more (less) opaque. Red line points to the preferred direction, and is projected onto the *XY* plane. **(E)** Same function as in **(D)** shown using a nested “matryoshkian” visualization of spheres of different radii (iso-*ω* shells).

Translations are straightforward to conceptualize and visualize, and it is equally straightforward to define functions on the space of translations. Unfortunately, however, rotations are far from straightforward, and many representations have been devised – such as Euler angles and axis–angle representations – to help scientists and engineers visualize and make sense of them. The challenge is that representations such as Euler angles necessarily have coordinate singularities, which complicate the definition of functions over the full rotation space. Unit quaternions provide a singularity-free alternative: the configuration of SO(3) can be expressed as the antipodal quotient S^3^*/*(**q** ∼ ™**q**), in which diametrically opposite points, **q** and −**q**, are identified. Here, **q** denotes a unit quaternion: a singularity-free unit vector in ℝ^4^ that represents a rotation. Unit quaternions form S^3^, which is a double cover of SO(3), because **q** and −**q** correspond to the same rotation matrix (SI Appendix, Eq. **??**).

The manifold describing 3D pose is called the Special Euclidean Group, SE(3). As a manifold, SE(3) is the Cartesian product of ℝ^3^ and SO(3), namely SE(3) = ℝ^3^ × SO(3). This product structure means that for every point in ℝ^3^ representing a position, there is a copy of SO(3) representing all orientations. Each element of SE(3) corresponds to a unique 3D pose specified by an ordered pair *g* = (**p**, *R*). This structure naturally lends itself to testing whether neural tuning functions are multiplicatively separable. Consistent with that possibility, separability increases along the visual hierarchy computing 3D object pose (Doudlah et al., 2022). Furthermore, deviations from separability are associated with lower-level representations in the visual system (Rosenberg et al., 2023). Likewise, head-direction selectivity and positional representations are largely separable in hippocampal and parahippocampal networks (Cacucci et al., 2004; Hardcastle et al., 2017). These higher-level separable representations of position and orientation in the limbic system must also be anchored to the external world, potentially through coupling between visually anchored representations in area 29e and path-integration circuits in medial entorhinal cortex (Secer et al., 2026). The positional component of these representations in ℝ^*N*^ (i.e., place cells (O’Keefe, 1976)) is derived in part from input from grid cells, although their population activity within individual modules evolves on a toroidal manifold rather than reflecting the topology of the physical space they encode (Gardner et al., 2022). Below, we focus on the nontrivial geometry of SO(3). Under a separability assumption, position can be incorporated by multiplying the orientation-dependent function by a function on ℝ^*N*^, such as a Gaussian or sigmoid over **p** (see Discussion).

### Neuronal encoding models on SO(3)

If neural systems encode 3D orientation consistent with S^3^*/*(**x** ∼ −**x**), then neuronal activity should be described by functions over that space. We therefore next derive two such functions, which are complementary in that one operates on quaternions and the other operates on rotation matrices.

In the field of geomagnetism, a standard model for dispersion data is the Bingham function, an antipodally symmetric function on S^2^ (Bingham, 1974). Here we extend that model to S^3^ to describe 3 DOF orientation.

The Bingham function, Ψ : S^3^ *→* ℝ^1^, maps points on the 3-sphere to the real line:

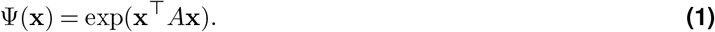

Here, **x** = [*w, x, y, z*]^*⊤*^ is a unit quaternion, and *A* = *A*^*⊤*^ ∈ ℝ^4*×*4^ is a symmetric matrix. Let

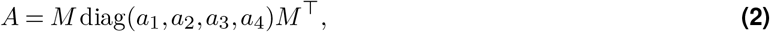

where the columns *µ*_*i*_ of *M* are orthonormal eigenvectors of *A* and *a*_*i*_ are the corresponding eigenvalues. We label the eigenvalue-eigenvector pairs such that *a*_4_ ≥ *a*_3_ ≥ *a*_2_ ≥ *a*_1_. The eigenvector matrix *M* is orthogonal.

Defining **y** = *M*^*⊤*^**x** such that 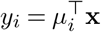 gives

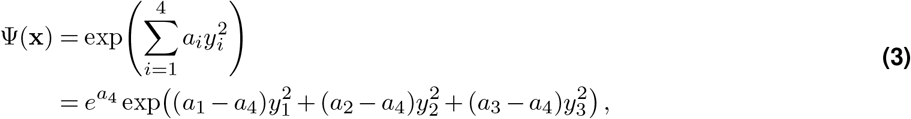

where the second equality follows from the fact that 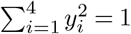. Define *λ*_*i*_ := *a*_*i*_ *−a*_4_ *≤* 0 for *i* = 1, 2, 3. Introducing offset (*dc*) and gain (*G*) terms, the Bingham tuning function on S^3^ becomes

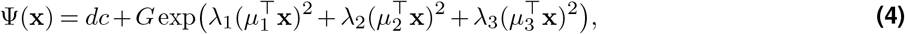

where the constant factor *e*^*a*4^ is absorbed into *G*. Because *λ*_*i*_ ≤ 0 and *G >* 0, the global maxima occur at **x** = ±*µ*_4_. Since antipodal unit quaternions represent the same rotation, *µ*_4_ specifies the preferred orientation.

What remains is to parameterize *M*. We define the Euler-angle rotation sequence as (1) roll about the *x*-axis (−180° ≤ *ϕ <* 180°), (2) pitch about the *y*-axis (−90° ≤ *θ* ≤ 90°), and (3) yaw about the *z*-axis (0° *ψ <* 360°), with reference orientation (*ϕ*_ref_, *θ*_ref_, *ψ*_ref_) = (0°, 0°, 0°), which corresponds to the identity quaternion **x**_ref_ = [1, 0, 0, 0]^*⊤*^.

The preferred orientation (*ϕ*^*\**^, *θ*^*\**^, *ψ*^*\**^) corresponds to the quaternion **x**^*\**^ = [*w*^*\**^, *x*^*\**^, *y*^*\**^, *z*^*\**^]^*⊤*^ (SI Appendix, Eq. **??**). Left quaternion multiplication, used to rotate the basis quaternions *i* = [0, 1, 0, 0]^*⊤*^, *j* = [0, 0, 1, 0]^*⊤*^, and *k* = [0, 0, 0, 1]^*⊤*^ by **x**^*\**^, defines a 4 *×* 4 orthonormal matrix:

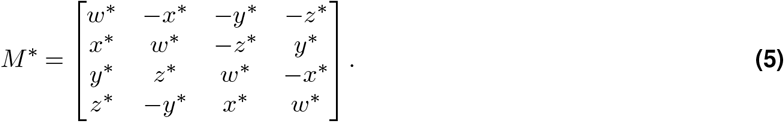

Its first column is **x**^*\**^, and its remaining columns form an orthonormal basis for the tangent space of S^3^ at **x**^*\**^.

The three eigenvectors orthogonal to *µ*_4_ = **x**^*\**^ determine the principal directions along which firing rate falls off from the preferred orientation. Their orientation within the tangent space is specified by a three-DOF rotation,

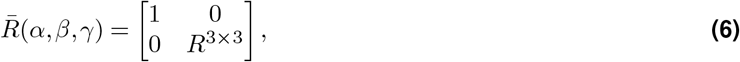

where *R*^3*×*3^ = *R*_*z*_(*γ*)*R*_*y*_(*β*)*R*_*x*_(*α*). We denote 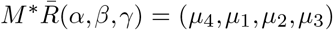. To restore the convention that column *i* of *M* is *µ*_*i*_, define the column-permutation matrix

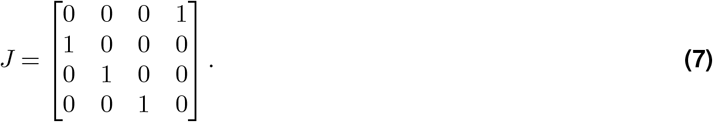

This results in 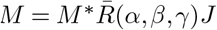 (SI Appendix, Eqs. **??, ??**). In total, the Bingham on S^3^ has 11 parameters: 3 for preference (*ϕ*^*\**^, *θ*^*\**^, *ψ*^*\**^), 6 for shape (*α, β, γ, λ*_1_, *λ*_2_, *λ*_3_), an offset (*dc*), and gain (*G*).

Since the quaternion double cover S^3^ lies in 4D Euclidean space, it cannot be directly visualized. However, neuronal activity over all three rotational DOFs can be projected into a solid sphere in ℝ^3^ (Fig. 2C). The solid sphere coordinates are an axis-angle representation in which 3D orientation is described by an axis (*u*; a unit 3-vector) and angle (0 ≤ *ω* ≤ *π* radians). The solid sphere has radius *π*, and each point has the coordinate *ω* · *u* (Fig. 2C, blue). For a given rotation axis, increasing the rotation angle moves a point further from the center. A rotation of *π* about *u* corresponds to the same orientation as *π* about −*u* (Fig. 2C, pink). Each point on the outer shell is therefore “glued” to its antipodal point. Geometric depictions of the Bingham on S^3^ are shown in Fig. 2D, E. Below, this method is used for the first time to visualize conjunctive neuronal tuning for roll, pitch, and yaw.

The field of robotics provides a complementary function, the Chillingworth-Marsden-Wan-Koditschek (CMWK) function (Chillingworth et al., 1982; Koditschek, 1988), for kinematic planning and control. The CMWK operates on rotation matrices, and is defined as *φ*(*R*) = Tr[*P R*], where *P* ∈ ℝ^3*×*3^ is a symmetric matrix with all the pairwise sums of the eigenvalues being nonnegative (SI Appendix), *R* ∈ ℝ^3*×*3^ is a rotation matrix, and Tr is the trace operator. To model neuronal tuning, we replace *R* with the rotational error 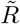 between the current orientation *R* and the preferred orientation *R*^*\**^, and exponentiate *φ*(*R*) and introduce offset (*dc*) and gain (*G*) terms:

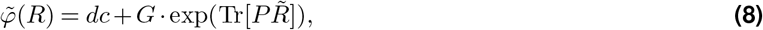

where 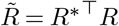. Here, *R*^*\**^(*ϕ*^*\**^, *θ*^*\**^, *ψ*^*\**^) = *R*_*z*_(*ψ*^*\**^)*R*_*y*_(*θ*^*\**^)*R*_*x*_(*ϕ*^*\**^) and *R*(*ϕ, θ, ψ*) = *R*_*z*_(*ψ*)*R*_*y*_(*θ*)*R*_*x*_(*ϕ*). *P* is defined as:

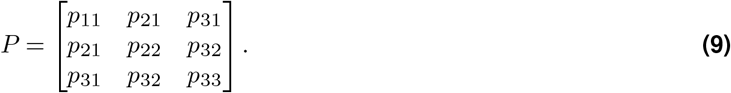

The CMWK thus has 11 parameters: 3 in 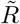 for the preference, 6 in *P* to set the shape, an offset (*dc*), and gain (*G*). In particular, the eigenvalues and eigenvectors of *P* set the tuning bandwidths and directions (SI Appendix).

In the SI Appendix (mathematical derivations) we prove the equivalency of the Bingham and CMWK on S^3^. For describing tuning functions on this manifold, the Bingham is preferable because the parameters are directly interpretable as familiar properties (e.g., bandwidths). Movement constraints can restrict SO(3) to submanifolds, while object symmetries identify indistinguishable orientations to form quotient manifolds. Although the Bingham and CMWK are equivalent when restricted to the same subset of SO(3), their natural parameterizations may differ in suitability for these new domains. Below, we show how the resulting manifold geometry and symmetry caused by object shape and movement constraints determine which function is appropriate.

### Spatial experience is constrained along manifold taxonomies

The 3D experience of a species depends on anatomical and physiological factors as well as its ecological niche (Rosenberg et al., 2025). For instance, binocular vision facilitates depth perception but limits the field of view. Visual experience is also shaped by an object’s features and mobility. Terrain and mode of locomotion similarly constrain navigation circuitry. Because all neural systems are subjected to constraints, neural coding will generally occur over manifolds derived from SE(3), which we next organize along manifold taxonomies (Fig. 3). Within a taxonomy, the most superordinate manifold describes the general encoding problem and each subsequent subordinate manifold describes a more constrained context. In this section, we first describe a *universal manifold taxonomy* describing 6 DOF pose. We then illustrate how visual symmetries and shape also transform the manifold geometries along a taxonomy of perceptibility, showing how the resulting manifold geometries yield testable predictions about neuronal coding.

**Figure 3.**
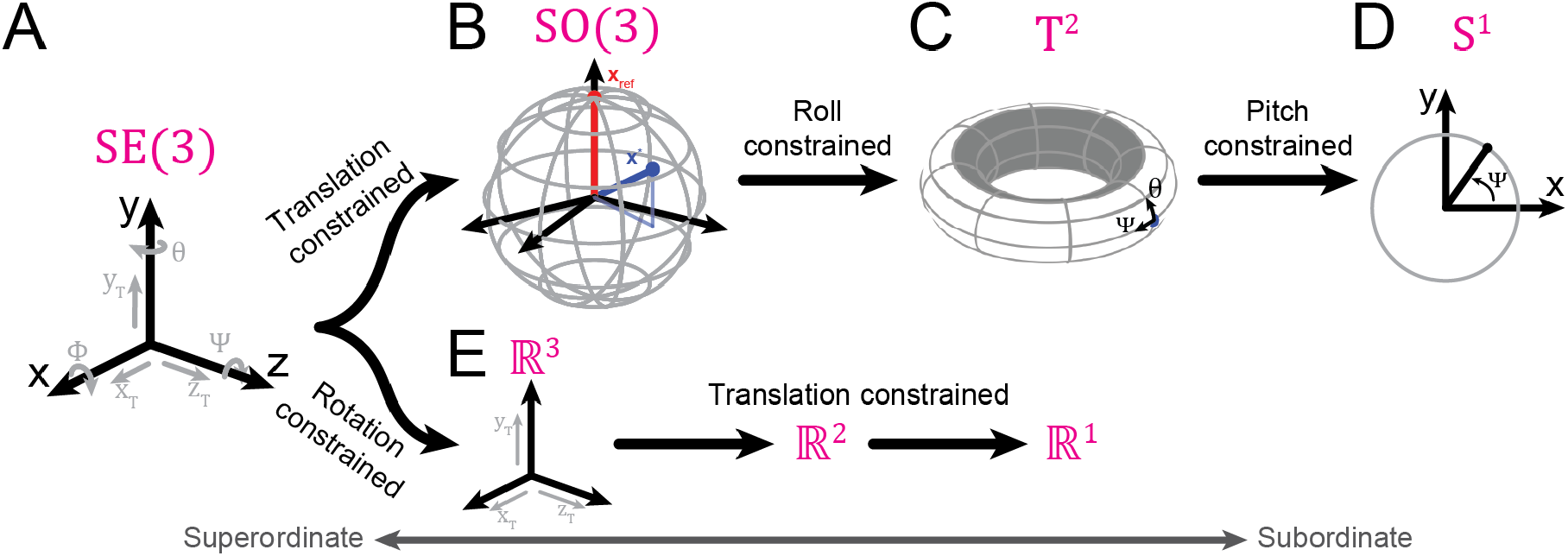
Universal manifold taxonomy. **(A)** The superordinate manifold, SE(3), describes the three translational (*x*_*t*_, *y*_*t*_, *z*_*t*_) and three rotational, roll (*ϕ*), pitch (*θ*), and yaw (*ψ*), DOFs of 3D experience. Subordinate manifolds diverge depending on the constrained DOFs. **(B)** Along a translation constrained branch, the subordinate manifold is SO(3), describing 3 DOF rotations. **(C)** Constraining roll, the manifold becomes T^2^, describing pitch (*θ*) and yaw (*ψ*). **(D)** If pitch is further constrained, the resulting manifold is relegated to S^1^, the most subordinate structure along this branch. **(E)** Along a rotation constrained branch, SE(3) is reduced to ℝ^3^, describing the 3 DOF translations. Subsequent translational constraints transform the manifold to ℝ^2^ and then ℝ^1^.

#### Universal manifold taxonomy for movement

The most superordinate manifold describing spatial experience is SE(3), containing the 6 DOFs for translations and rotations (Fig. 3A). The taxonomic structure then diverges depending on which DOFs are restricted. Along a branch that is “translation free”, the first subordinate manifold describes 3 DOF rotations, SO(3) (Fig. 3B). As described above, the geometric structure is S^3^*/*(**x** ∼ −**x**), commonly denoted ℝ ℙ^3^. If roll rotations are restricted, i.e., fixed, the resulting subordinate manifold is a torus, T^2^, if pitch is extended to −180° *< θ* ≤ 180° to capture all possible roll-free orientations (Fig. 3C). Thus, pitch and yaw are separable rotations forming the minor and major circles of a torus, respectively. If pitch rotations are then restricted, the most subordinate manifold along this branch is S^1^, describing yaw rotations (Fig. 3D). A “rotation-free” branch can similarly be defined, starting with ℝ^3^ describing the 3 DOF translations (Fig. 3E). As translational DOFs are restricted, the subordinate manifolds become ℝ^2^ and then ℝ^1^.

This illustrated manifold taxonomy separates translation- and rotation-free branches. However, subordinate manifolds of SE(3) are not restricted to a single type of movement. When translational and rotational constraints are independent, mixed constraints yield Cartesian-product manifolds. For instance, if roll rotations are restricted and translations are limited to 1 DOF, the resulting subordinate manifold would be T^2^ × ℝ^1^. More generally, constraints may couple translation and rotation, in which case the resulting configuration space need not be a Cartesian product of separately defined translational and rotational manifolds. Under the assumption of independent constraints, the manifolds in Fig. 3 can be combined to identify other subordinate structures of SE(3).

#### Illustrative case 1: Manifold taxonomy of perceptibility for a viewed bar

Visual and motor function are often studied using oriented stimuli such as bars, which we use here to introduce how visual symmetries and object shape transform a manifold’s geometry. This deductive process reveals how seemingly innocuous changes in experimental design can have nontrivial implications for the predictions for neuronal coding.

We proceed by first defining a coordinate frame and reference orientation for a viewed bar. Consistent with common practice in visual neuroscience, we use a left-handed coordinate frame and define the bar’s reference orientation to be along the *x*-axis, (*ϕ*_ref_, *θ*_ref_, *ψ*_ref_) = (0°, 0°, 0°) (Fig. 2A).

For a texture-rich bar, the most superordinate manifold is SE(3), describing the 6 DOFs for rotations and translations. To investigate how visual symmetries transform the manifold’s geometry, we focus on rotational DOFs in this section, yielding the subordinate manifold SO(3) (Fig. 4A). If neural systems encode spatial experience consistent with the geometry of SE(3), then neuronal representations may be described by functions that operate over the manifold or subordinate manifolds of SE(3). Thus, for SO(3), population activity encoding orientation can be modeled by functions that operate on S^3^*/*(**x** *∼ −***x**). As such, the Bingham function on S^3^ is a candidate model for describing the responses of orientation-selective neurons.

**Figure 4.**
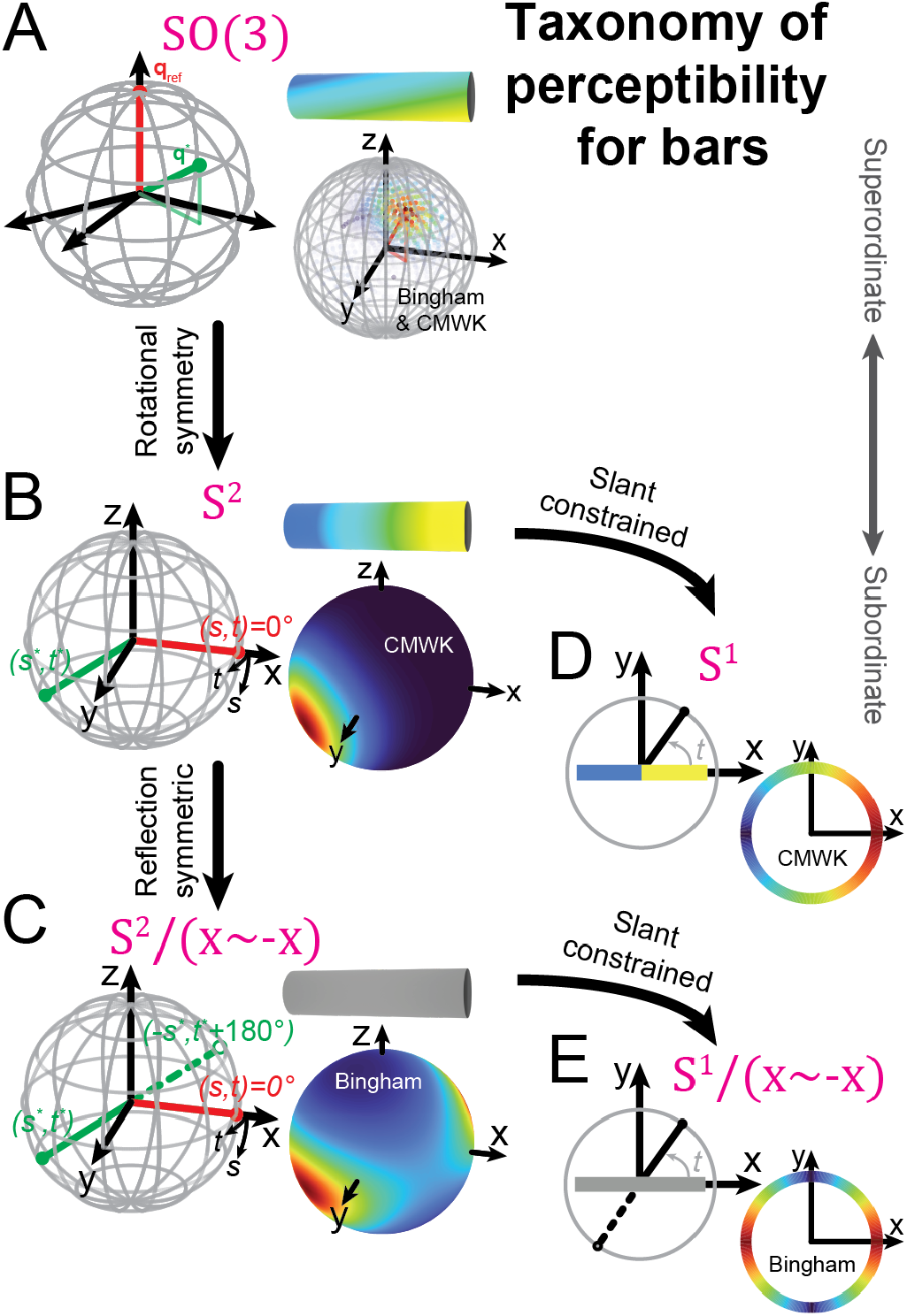
Manifold taxonomy of perceptibility for bars. **(A)** For a texture-rich bar, the superordinate manifold describing rotations is SO(3). This geometry predicts that neuronal responses will be described by Bingham/CMWK functions, visualized using the solid sphere projection. **(B)** A bar with rotationally symmetric texture results in the subordinate manifold S^2^, predicting that neuronal responses will be described by the CMWK function. **(C)** An “achiral” bar, such as one lacking any texture, has the subordinate manifold S^2^*/*(**x** ∼ −**x**), predicting that tuning functions will be described by Bingham functions. **(D)** If slant rotations are constrained for a rotationally symmetric bar, the resulting manifold is S^1^, predicting that neuronal responses will be described by the CMWK function. **(E)** If slant is constrained for an achiral bar, the manifold reduces to S^1^*/*(**x** *∼ −***x**), predicting that neuronal responses will be described by Bingham functions.

Proceeding from SO(3), visual symmetry constraints define a taxonomy of perceptibility. Consider if the bar’s texture is changed such that rotations about the long axis (roll) are not discernible (Fig. 4B). The kinematics allow for roll, but neurons will be insensitive to that rotation, resulting in orientation metamers. In effect, the orientation of a bar with a roll-free texture is fully described by the direction of a unit vector in ℝ^3^. The corresponding manifold is therefore S^2^, a subordinate manifold of SO(3). Because opposite points on S^2^ correspond to distinct bar orientations, a Bingham function defined directly on S^2^ would impose an unwanted antipodal symmetry. We therefore use the CMWK function on SO(3), or equivalently a Bingham function on unit quaternions, parameterized to be invariant to roll. For the CMWK, this is achieved by defining

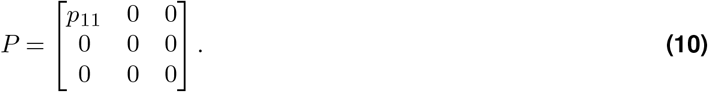

To align the parameterization of the CMWK on S^2^ with vision-science terminology (Stevens, 1983), we refer to pitch as slant and yaw as tilt. For bars, slants of ±90° result in self-occlusion, creating a perceptual singularity. Furthermore, yaw rotations at these slants become indistinguishable because they correspond to rotations about the bar’s long axis. We therefore restrict slant to *−*90° *< s <* 90°, thereby excluding the north and south poles of S^2^ (Fig. 4B, green circles). Tilt remains unrestricted, 0° *≤ t <* 360°. Thus, slant and tilt correspond to the elevation and azimuth coordinates on S^2^, respectively. The rotation matrix specifying the orientation of a roll-free bar is then

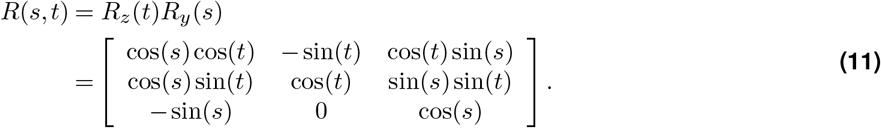

The CMWK is therefore parameterized by *P* (Eq. 10) and 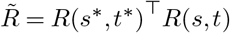 (Eq. 11), where *s*^*\**^ and *t*^*\**^ are the slant and tilt preferences.

Now consider a bar with a roll-free, reflection-symmetric texture (Fig. 4C). Because the two ends of this “achiral” bar are indistinguishable, the slant–tilt configurations (*s, t*) and (−*s, t* + 180°) are perceptually equivalent. On S^2^, these configurations map to antipodal points, **x** and −**x**. Therefore, the predicted manifold is relegated to S^2^*/*(**x** ∼ −**x**). This topology explains why an achiral bar’s tilt is *π* periodic at *s* = 0° but 2*π* periodic at non-zero slants. In this case, the manifold’s antipodal symmetry predicts that single-neuron responses will be described by a Bingham on S^2^ (SI Appendix), which naturally captures the slant-dependent periodicity of tilt:

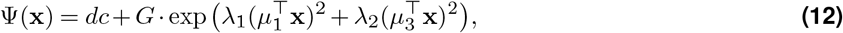

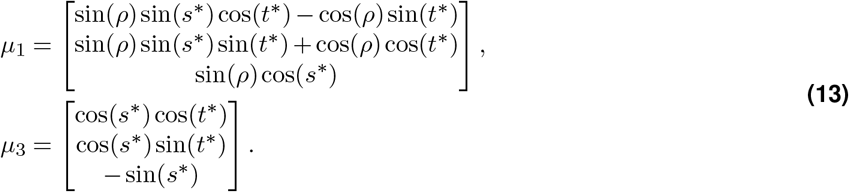

Here, *λ*_1_ ≤ 0 determines the tuning anisotropy, *λ*_2_ ≥ 0 sets the bandwidth, and *ρ* rotates Ψ(**x**) about (*s*^*\**^, *t*^*\**^) (coupling *s* and *t*). Note that *µ*_3_ is the coordinate mapping to S^2^*/*(**x** ∼ −**x**).

Further constraining a rotationally symmetric bar to the frontoparallel plane (i.e., fixed slant, *s* = 0°) reduces the predicted manifold to S^1^ (Fig. 4D), with orientation parameterized as:

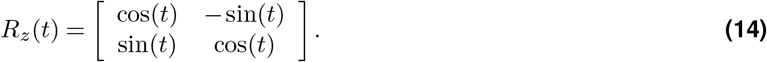

In this case, orientation is 2*π* periodic because the bar’s ends are discernible, predicting that the CMWK is suitable for describing neuronal tuning. Now 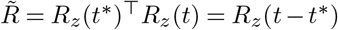, and *P* is a diagonal matrix since off-diagonal elements cancel after 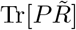:

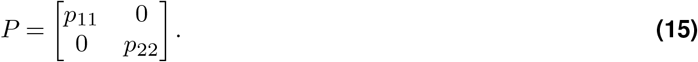

Substituting 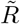 and *P* into Eq. 8 yields:

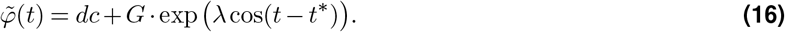

Here, *λ* = *p*_11_ + *p*_22_. Thus, the CMWK reduces to a 2*π*-periodic von Mises function in 1D.

However, for a textureless bar restricted to the frontoparallel plane, the previous S^1^ manifold is relegated to S^1^*/*(**x** ∼ −**x**) (Fig. 4E). Orientation is then *π* periodic, corresponding to stimuli used to study V1 (Hubel and Wiesel, 1968). Accordingly, neuronal tuning can be described by a Bingham function on S^1^—equivalently, a *π*-periodic von Mises tuning function (SI Appendix):

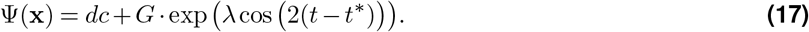

#### Illustrative case 2: Testing deductive manifold predictions

Here we use the visual encoding of 3D planar surface orientation to test if the deductive framework yields manifolds that predict the response profiles of single neurons and latent population activity. Manifold taxonomies for planes largely parallel those for bars, and we go straight to an “achiral plane” for which roll (rotation about the surface normal) and the two sides are not discernible. For achiral planar surfaces, 3D orientation is fully described by a unit 3-vector, the plane’s normal (Fig. 5A, red arrow), yielding the manifold S^2^*/*(**x** ∼ −**x**) (Fig. 5B).

**Figure 5.**
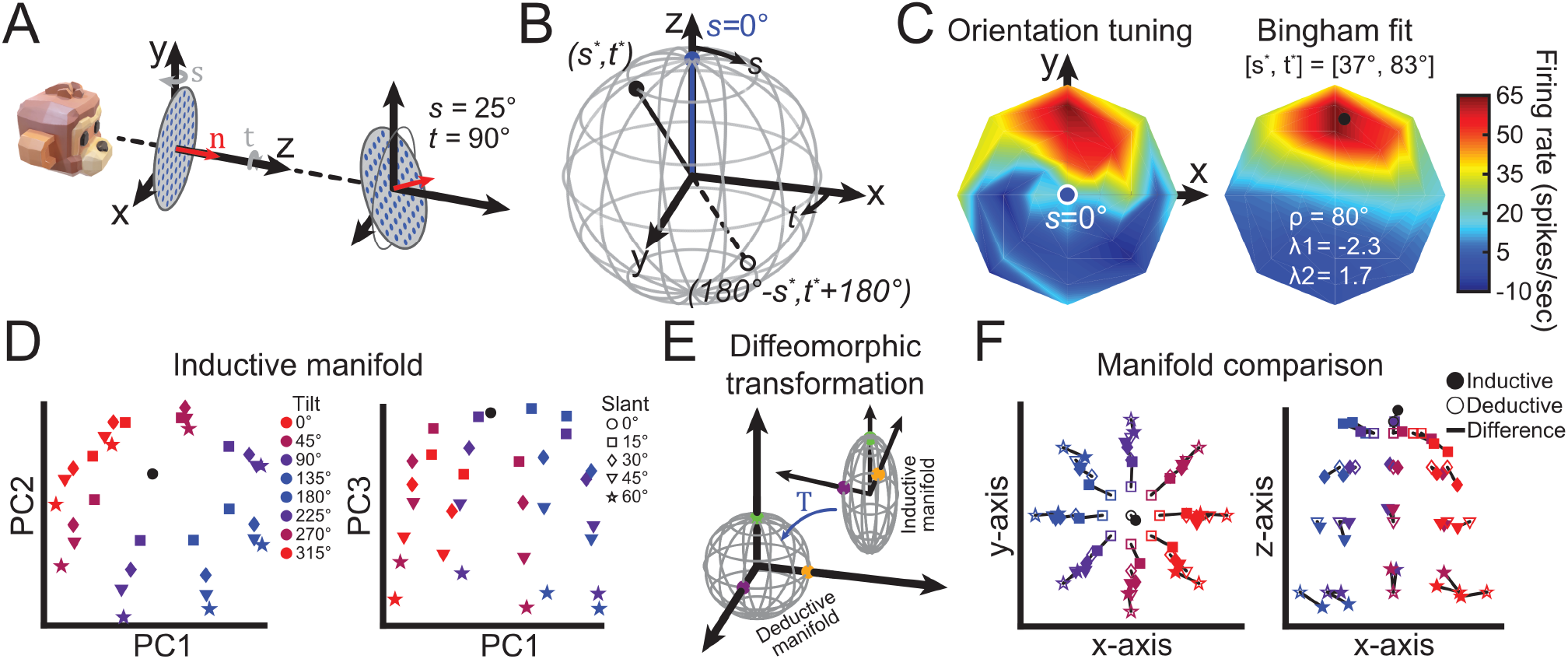
Manifold for planes and comparison of deductive and inductive approaches. **(A)** Coordinate frame for the 3D orientation of planar surfaces. The reference orientation is frontoparallel (left), defined by the surface normal (*n*, red arrow) pointing along the *z*-axis. Slant (*s*) and tilt (*t*) are rotations about the *y*- and *z*-axes, respectively. *Right:* An oriented surface is shown (*s* = 25°, *t* = 90°). Grey circle depicts the reference orientation to aid visualization. **(B)** The manifold geometry is S^2^*/*(**x** *∼ −***x**), where *s* and *t* are mapped to the colatitude and azimuth, respectively, with the reference orientation mapped to the north pole (blue). **(C)** Example baseline-subtracted responses from CIP (left) with Bingham fit (right). Data are visualized using an equatorial projection. Preferred *s* and *t* values are indicated on the Bingham fit (black circle). Red (blue) hues indicate higher (lower) firing rates. **(D)** Inductive manifold derived using PCA. Projections are shown for PC1,2 (left) and PC1,3 (right). Orientation is coded separately for *t* (colors) and *s* (shapes). **(E)** Schematic illustrating the method to account for the fitted affine transformation (*T*, blue arrow), a local change of coordinates between the inductive and deductive manifolds. Colored points are shown to visualize the transformation between manifold shapes. **(F)** Comparison of manifold geometries derived using inductive (filled shapes) and deductive (open shapes) approaches. PCA data in **(D)** are projected onto the deductive manifold after applying the coordinate transformation. Black lines indicate differences. Color and shapes are consistent with **(D)**.

A different coordinate mapping is required to capture the perceptual and coordinate singularities of planes than for bars. The frontoparallel plane (*s* = 0°) is used as the reference orientation. Slant and tilt map to colatitude and azimuth, but *s* = 0°, located at the pole, results in a perceptual singularity since tilt rotations are not discernible, and 90°, located at the equator, leads to self-occlusion. The different mappings reflect that tilt is defined for a bar in the frontoparallel plane (the equator, Fig. 4B), but is undefined for a frontoparallel plane itself (the pole, Fig. 5B, blue). The manifold geometry again predicts the Bingham will describe neuronal tuning, but a reparameterization is needed to account for the coordinate mapping (*µ*_3_) (Rosenberg et al., 2013):

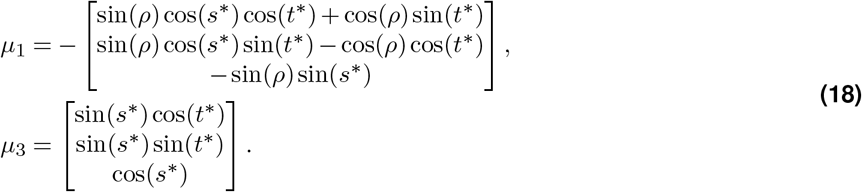

In macaques, 3D object pose is encoded in the caudal intraparietal (CIP) area (Rosenberg et al., 2023). The orientation selectivity of a CIP neuron for an achiral plane is shown in Fig. 5C (left). Consistent with the predictions, the responses are well described by a Bingham on S^2^*/*(**x** ∼ −**x**) (right). Indeed, across 380 neurons (Chang et al., 2020), the mean fit was *r* = 0.87 ± 0.005 standard error (SI Appendix, detailed methods). Given the strong model performance and since topological validity does not imply physiological relevance, a natural question is whether all of the parameters are necessary. For instance, beyond key parameters (e.g., preference, bandwidth), the necessity of others such as coupling terms (*ρ*) is less apparent. We therefore performed a model comparison and found that tuning was best described with *ρ* as a free parameter (SI Appendix, Fig. **??**). A productive feature of the deductive framework is the efficiency of model identification, but this comparison highlights that model refinement requires empirical tests to define the most parsimonious characterization of neural data.

We next test if the deductive manifold S^2^*/*(**x** ∼ −**x**) describes the latent population activity of CIP using principal component analysis (PCA) and an affine least-squares alignment of the neural and predicted coordinates. Indeed, direct comparisons of deductive and inductively inferred manifolds are needed to test the framework since there is no *a priori* requirement for neural systems to conform to the inherent geometries of encoding problems.

The first three principal components (PCs) explained 64% of the variance in the population data. Consistent with a roughly top-down view of the deductive manifold, a plot of PCs 1 and 2 revealed a spoke-like structure with tilt organized as the angular variable and slant as the radial variable, with the frontoparallel plane (*s* = 0°) near the center (Fig. 5D, left; black circle). Consistent with a side-view of the deductive manifold, a plot of PCs 1 and 3 likewise revealed that slant was organized along layers similar to lines of latitude (Fig. 5D, right).

To quantify agreement between the inductively derived neural manifold and the deductively predicted geometry, we fit an affine mapping between the sampled neural-response patch and its corresponding predicted orientation patch, minimizing their discrepancy (SI Appendix, Detailed Methods, Eq. **??**). Although the sampled slants, 0°–60°, with all tilts form a disk-like patch rather than the full projective plane, the mapping provides a quantitative test of local geometric agreement. For intuition, an ellipsoid can be smoothly deformed into a sphere, illustrating equivalence under a diffeomorphism (Fig. 5E). Applied to the sampled orientations, the transformation yielded low residual error and close correspondence between the neural and predicted patches (Fig. 5F; SI Appendix). This finding supports agreement between the neural and predicted geometries over the sampled range of surface orientations.

#### Illustrative case 3: Head direction manifolds

A high-fidelity representation of head direction is critical for navigation (Taube et al., 1990; Taube, 1995; Finkelstein et al., 2015; Angelaki et al., 2020), postural control (Angelaki and Cullen, 2008), gaze stabilization (Leigh and Zee, 2015), and balance (Horak, 2006). Head direction has 3 DOFs, but most previous work focused on lower-dimensional subspaces. We next show how the iterative relaxation of movement constraints transforms the deductive head direction manifold from a 1 DOF compass to the full dimensionality of SO(3). In parallel, we establish topologically correct, expressive functions describing neuronal responses in bats, mice, and macaques.

##### 1 DOF head direction

We first briefly consider head direction in the *xy* plane. Since planar orientation has one DOF, the deductive manifold is S^1^ and the CMWK function with *P* a 2 × 2 diagonal matrix and *R*(*ψ*)^2*×*2^ is topologically correct. In this case, the CMWK reduces to a 2*π*-periodic von Mises function (Eq. 16). This function is widely used to quantify head direction tuning (Taube et al., 1990; Skaggs et al., 1994; Taube, 1995; Secer et al., 2025), providing empirical support that spatial experience is encoded on subordinate manifolds of SO(3) when movement is constrained.

##### 2 DOF head direction

A seminal study of 3D head direction selectivity examined pitch–yaw tuning in the dorsal presubiculum of Egyptian fruit bats (Finkelstein et al., 2015). Comparisons of neuronal activity in upright and inverted poses led to the inductive inference that pitch and yaw are encoded on a toroidal manifold. In the Supplementary Information (mathematical derivations), we provide a formal proof that the manifold for roll-free head direction is indeed a torus by constructing a diffeomorphism between T^2^ and the set of rotation matrices describing roll-free head direction. Intuitively, this arises because pitch (*θ*) and yaw (*ψ*) provide a singularity-free parameterization in which both variables span 360°, yielding S^1^ *×* S^1^ = T^2^ (Fig. 6A).

**Figure 6.**
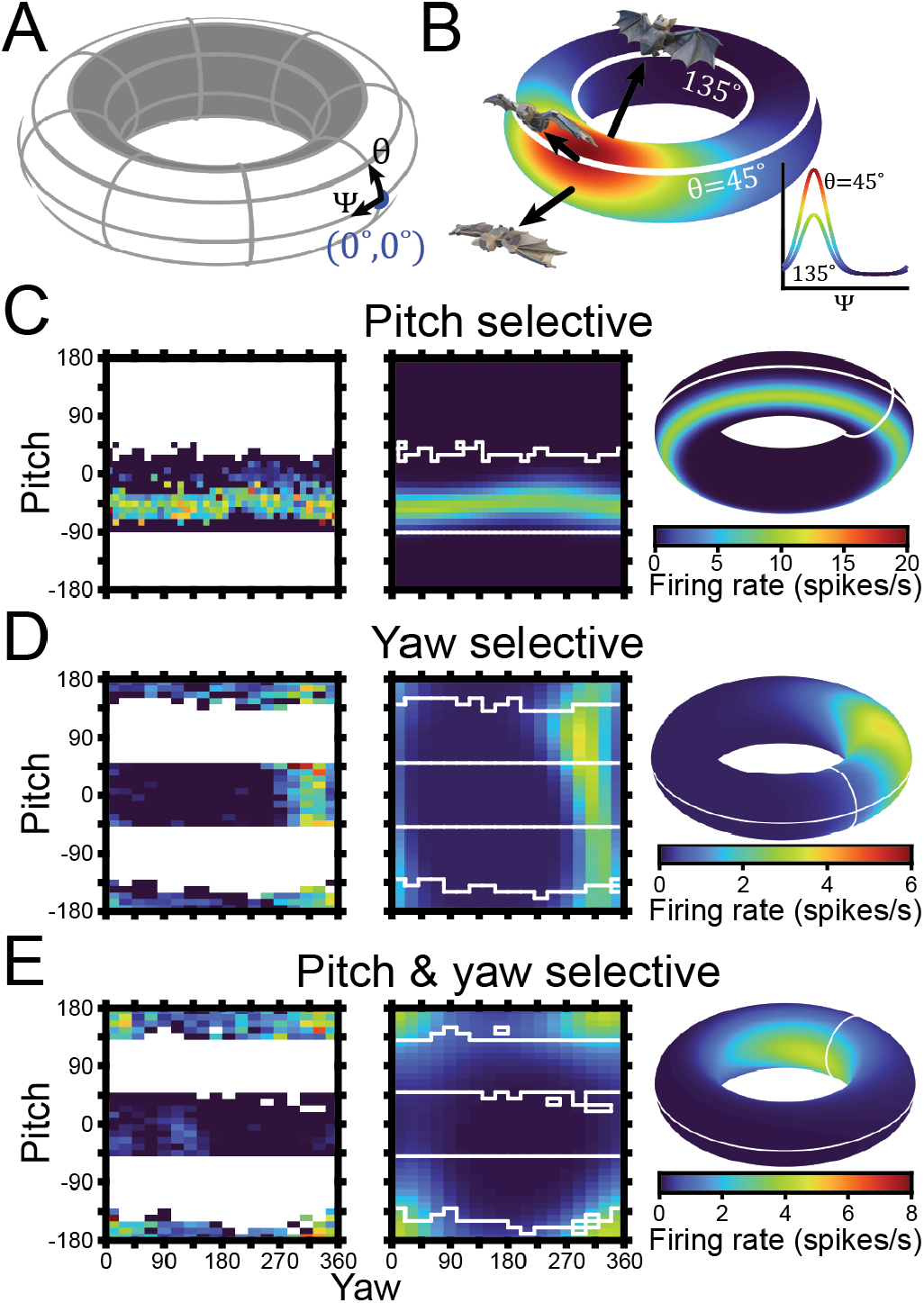
Toroidal manifold of roll-free head orientation. **(A)** Toroidal manifold describing pitch (*θ*) and yaw (*ψ*). The reference orientation (*θ, ψ*) = (0°, 0°) is marked by a blue point. **(B)** Illustrative CMWK function. Black arrows mark head direction at three pitches. White iso-yaw lines correspond to the yaw tuning curves in the inset. **(C)-(E)** Head direction tuning curves. *Left:* Neuronal responses. White space indicates unsampled head directions. *Middle:* CMWK fits. White lines demarcate sampled regions. *Right:* CMWK fits on the torus. White circles mark *θ* = 0° and *ψ* = 0°. Red (blue) hues indicate higher (lower) firing rates. **(C)** Pitch selective neuron from the macaque anterior thalamus. **(D)** Yaw selective neuron from the bat dorsal presubiculum. **(E)** Joint pitch–yaw selective neuron from the bat.

To derive the CMWK function on T^2^, we follow the left-handed coordinate frame in Finkelstein et al. (2015) with the nose pointing in the +*x* direction and right ear in the +*y* direction. We then define *R*(*θ, ψ*), where pitch about the *y*-axis occurs first, *R*_*y*_(*θ*), followed by yaw about the *z*-axis, *R*_*z*_(*ψ*):

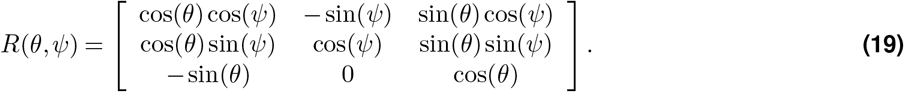

Setting 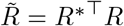, describes the two allowed rotational DOFs and sets the preferred orientation. Defining *P* as in Eq. 9 to determine the function’s expressiveness, the CMWK on T^2^ takes the same form as Eq. 8:

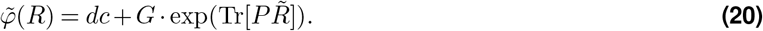

An example CMWK is shown on a torus in Figure 6B. A critical but perhaps nuanced distinction that has caused confusion is that an animal’s compass direction (azimuth) in the world is not the same as its angular head direction (yaw) as an Euler parameterization. To illustrate an important implication, yaw tunings at two pitches (*θ* = 45°, upright; 135°, inverted) are shown in the inset. Although the compass direction in which the bat is pointed naturally shifts by 180° when it inverts, the preferred yaw remains the same. Put succinctly, if pitch causes an animal’s pose to invert, its yaw (head direction) remains the same but the azimuth (defined as the compass direction in which the nose is pointing in the world, which is undefined at *θ* = *±π/*2) jumps by 180°:

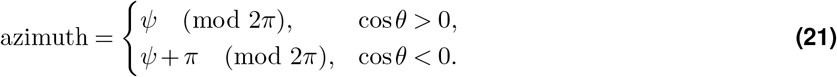

To demonstrate the expressive capacity of Eq. 20 for describing head-direction tuning, we fit the model to responses from the macaque anterior thalamus (Supplementary Information, detailed methods) and bat dorsal presubiculum (Finkelstein et al., 2015). As predicted by the theory, strong cross-species correspondence between the CMWK on T^2^ and the activity of pitch-tuned (Fig. 6C), yaw-tuned (Fig. 6D), and joint pitch–yaw tuned (Fig. 6E) neurons was observed. These empirical observations further support that neuronal systems encode orientation on submanifolds of SO(3).

##### 3 DOF head direction

If all rotational constraints are eliminated, the manifold describing 3 DOF head direction is SO(3). Sampling the large combinatorial set of orientations across this space is a considerable mathematical and experimental challenge (Yershova and LaValle, 2004). Nevertheless, recent efforts inspired by work on how the head-direction reference frame is updated during 3D movement (Page et al., 2018) explored this selectivity using a motorized rotator to position mice at different orientations (Angelaki et al., 2020).

To test if the encoding of 3 DOF head direction is consistent with the geometry of SO(3), we fit the responses of two neurons from the study recorded in the cingulum fiber bundle (Fig. 7A,B; first column) with the Bingham function (Fig. 7; second column) (Supplementary Information, detailed methods). A nested shell visualization of the responses and fits is shown in Fig. **??**. For these fits, a right-handed coordinate frame is used, as in (Angelaki et al., 2020). Consistent with our theory that head direction is encoded on SO(3), the responses were well described by the Bingham function (A: *r* = 0.87; B: *r* = 0.76).

**Figure 7.**
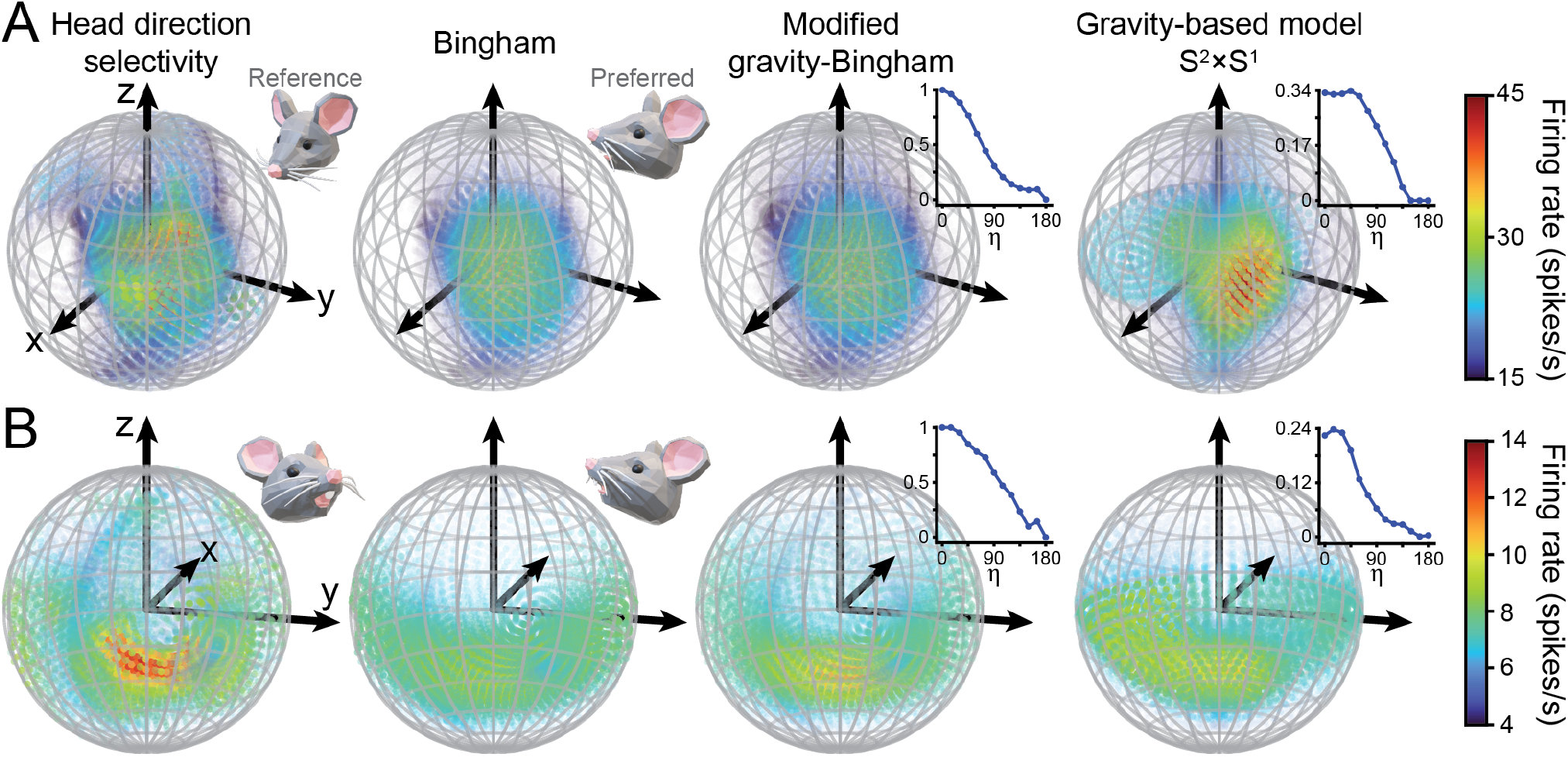
3 DOF head direction tuning. Responses of two neurons (**A, B**) recorded from the mouse cingulum fiber bundle for different head directions plotted using the axis–angle visualization (column 1). Model fits are shown for the Bingham function (column 2) with the corresponding preferred head direction shown (inset). The modified Bingham function (column 3) and gravity-based model (column 4) are also shown with corresponding head-tilt-dependent gain functions (insets). Red (blue) hues indicate higher (lower) firing rates.

Rather than on SO(3), the original study parameterized head direction as head tilt relative to gravity (a vector combining roll and pitch) and azimuth (Angelaki et al., 2020). That is, on the manifold S^2^ × S^1^. Correspondingly, head direction was quantified using a “gravity-based” model, the multiplication of head tilt and azimuthal functions (Laurens and Angelaki, 2019). Reflecting an observation that azimuthal tuning diminished and eventually became flat as the head tilted away from upright, the investigators incorporated a head-tilt-dependent gain function. To account for possible differences in neuronal tuning in postures less commonly adopted during natural behavior, such as full inversion, we modified the Bingham function to include the same gain function:

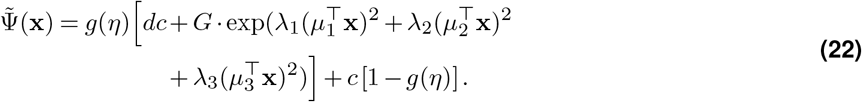

Here, *η* is the head tilt relative to gravity, with *η* = 0° corresponding to upright and *η* = 180° inverted. Following (Angelaki et al., 2020), *g*(*η*), bounded between 0 and 1, was defined using 11 free parameters sampled at 15° intervals with fixed ends *g*(0) = 1 and *g*(180) = 0, and *g*(*η*) was determined using linear interpolation in each interval. *c* sets the DC response at inverted postures. Modified gravity–Bingham fits are shown in Fig. 7 (third column). Despite the visual similarities, the modified version (A: *r* = 0.88; B: *r* = 0.81) yielded higher fit correlations and lower AIC values than the pure Bingham (A: AIC_modified_ = 8913.55, AIC_pure_ = 9093.28; B: AIC_modified_ = −2221.01, AIC_pure_ = −1424.57). This finding illustrates how empirical observations can be used to refine theory-derived tuning functions; here, we captured idiosyncratic features of neuronal activity in both commonly and rarely adopted postures.

Lastly, we compare the Bingham and gravity-based model fits. The gravity-based model fits are shown using the solid-sphere projection in Fig. 7 (last column). Although the gravity-based model performs well overall (A: *r* = 0.89; B: *r* = 0.79), it introduces features not present in the data, including an equatorial plane-like structure in Fig. 7A and a bowl-like structure in Fig. 7B. These features arise because of singularities in the model (see Supplementary Information, mathematical derivations) that do not exist for SO(3). The model also does not capture the arc-like structure in Fig. 7B, which is captured only by the gravity–Bingham. These results collectively support that the deductive framework provides a unified approach for exploring the neural basis of 3D spatial experience.

## Discussion

Neuronal coding theory is grounded in the manifold hypothesis (Fefferman et al., 2016), which implies that neural activity associated with 3D experiences lies near low-dimensional spaces embedded within ones of much higher dimension. We introduced a mathematical theory based on the premise that neural systems evolved to encode high-fidelity 3D spatial representation on latent submanifolds derived from SE(3), whose topology describes the six dimensions of translation (position) and rotation (orientation). We further outlined a deductive framework for applying the theory to empirical research. The approach defines manifold taxonomies along which object features and/or mobility are iteratively constrained. Formal characterization of the inherent manifold geometries makes explicit predictions for perception and neuronal selectivity. We focused on the nontrivial coding of orientation, but the geometries more generally correspond to well-recognized manifolds: (1) ℝ^2^ × S^1^*/*(**x** ∼ −**x**), the “Hubel and Wiesel” manifold for 2D position-invariant orientation tuning of V1 complex cells (Hubel and Wiesel, 1962), (2) ℝ^1^ × S^2^*/*(**x** ∼ −**x**) describing distance-invariant 3D orientation tuning in CIP for achiral planes (Chang et al., 2020), (3) ℝ^2^ × S^1^ describing 2D position-invariant planar head direction (Taube et al., 1990), (4) T^2^ describing roll-free head direction (Finkelstein et al., 2015), and (5) SO(3) describing 3 DOF head direction (Angelaki et al., 2020). These correspondences between deductive and inductive manifolds across species and encoding problems support the premise that neural systems evolved to encode spatial information over SE(3).

The first-principles approach outlined here is applicable to any encoding problem for which an inherent geometry amenable to geometric–topological analysis exists. In such cases, several benefits were shown. Methods were devised to concisely visualize conjunctive responses over 3 DOF rotations using a single, topologically correct depictive object. Data from the mouse cingulum fiber bundle highlighted how the global structure of neuronal responses can otherwise be masked, limiting interpretability and model comparison. The approach also guides the creation of topologically correct models for quantifying neuronal activity, regardless of dimensionality. Model selection can otherwise require substantial iterative effort (Swindale, 1998; Finkelstein et al., 2015; Angelaki et al., 2020). In that regard, the productiveness of the approach reflects that the properties of functions are constrained by manifold geometry, which it makes explicit.

The theory places the encoding of 3D visual object orientation and head direction within a common geometric framework derived from SO(3). This finding implies that if two systems utilize a common code with the same domain, then the manifolds describing the respective encoding problems must be diffeomorphic. At the same time, the emergence of non-diffeomorphic geometries, such as T^2^ and S^2^*/*(**x** ∼ −**x**), within and across manifold taxonomies had surprisingly deep ramifications for neuronal coding. A striking finding was that seemingly innocuous experimental design factors such as the use of bars or planes, or an object’s texture, fundamentally changed the encoding functions. Such changes could be related to why 3D feature selectivity in macaque V4 is largely independent of distance when assessed with bars (Hinkle and Connor, 2002) but not curved surfaces (Hegdé and Van Essen, 2005). Indeed, the factors determining a manifold’s geometry (and therefore constraining the neural code) are nuanced. Manifold taxonomies predict how neuronal response properties may flexibly change depending on context.

Empirical methods were shown to refine deductive findings in two ways. First, through the statistical assessment of the physiological relevance of topologically valid parameters. Second, species-dependent ecological and behavioral factors such as the orientation of the head relative to gravity (a critical signal for perceptual stability and postural control (Dakin and Rosenberg, 2018)) were shown to shape encoding functions. When the geometry of an encoding problem is complex or unknown, inductive approaches involving manifold inference (learning) may be most practical (Chang and Tsao, 2017; Yoon et al., 2024; Yao et al., 2026). Once inferred, however, the formal characterization of manifold geometries along taxonomies can provide deductive grounds for hypothesis testing. Still, inductive methods require large, unbiased datasets and are sensitive to data sparsity and noise (Helmer et al., 2016), potentially limiting the generality of findings. Moreover, because neural systems are not required to conform to the inherent geometry of an encoding problem, the comparison of deductive and inductive findings is valuable and discrepancies would be exciting because they could inform constraints shaping the evolution of neural systems.

The expressiveness of an encoding function depends not only on its critical points, but also on its free parameters and the variations they permit. Many of our tuning functions, including the unmodified Bingham and CMWK functions on SO(3), can describe unimodal tuning with the minimum number of nondegenerate critical points. Nevertheless, they provide highly flexible directional control within their respective parametric forms—their parameters control the principal curvatures around the preferred orientation, while coupling terms rotate these directions relative to the coordinate axes, enabling flexible anisotropic tuning but not arbitrary independent falloff in every direction. Certain object symmetries may instead produce multiple equivalent response modes when tuning is expressed over the full rotation space. Two ways to adapt the functions for such cases include: (1) harmonic expansions and (2) mixture distributions in which linear combinations of the same functional form are taken with fixed rotational offsets describing the symmetries. If modes beyond those required by object symmetry are observed in neuronal data, they could imply lower-level representations which indirectly covary with pose (Rosenberg et al., 2025) or conceivably the multiplexing of cognitive factors that associate multiple poses with a common behavioral contingency.

Geometric topology is a central but often implicit concept in neuroscience. At their core, tuning curves imply the existence of manifolds describing configuration spaces of encoded features. We showed that characterizing those spaces along taxonomies provides a deductive testbed for empirical research. In this work, the nontrivial geometry of rotations was emphasized. Further extensions will need to more deeply consider the joint representation of translations and rotations. We also focused on static pose, but note that the same manifolds can describe dynamic pose expressed as a parametric function of time. In that case, latent population activity traverses paths over the manifolds. Notwithstanding current limits, the framework shows that the study of inherent manifold geometries can reveal new insights into how neural systems enable animals to thrive in a complex and volatile world.

## Materials and Methods

For full details of the materials and methods, see Supplementary Information.

## Data availability

Previously published bat and mouse head direction data were provided with permissions from the original authors.

## Acknowledgments

We thank Dora Angelaki, Arseny Finkelstein, Jean Laurens, and Nachum Ulanovsky for sharing head direction cell data. This work was supported by NIH grant NS128586 (B.K. and A.R.), Retina Research Foundation Edwin and Dorothy Gamewell Professorship at the McPherson Eye Research Institute and NIH grants EY029438 and EY035005 (A.R.), as well as AFOSR FA9550-23-1-0722 and NIH grant U01NS131438 (N.J.C.). Further support was provided by NIH grant P51OD011106 to the Wisconsin National Primate Research Center.

## References

Angelaki, D. E. and Cullen, K. E. (2008). Vestibular system: the many facets of a multimodal sense. Annu. Rev. Neurosci., 31(1): 125–150.

Angelaki, D. E., Ng, J., Abrego, A. M., Cham, H. X., Asprodini, E. K., Dickman, J. D., and Laurens, J. (2020). A gravity-based three-dimensional compass in the mouse brain. Nature communications, 11(1):1855.

Bingham, C. (1974). An antipodally symmetric distribution on the sphere. The Annals of Statistics, 2(6):pp. 1201–1225.

Bonhoeffer, T. and Grinvald, A. (1991). Iso-orientation domains in cat visual cortex are arranged in pinwheel-like patterns. Nature,353(6343):429–431. doi: 10.1038/353429a0.

Bressloff, P. C. and Cowan, J. D. (2003). A spherical model for orientation and spatial-frequency tuning in a cortical hypercolumn.Philos Trans R Soc Lond B Biol Sci, 358(1438):1643–1667. doi: 10.1098/rstb.2002.1109.

Cacucci, F., Lever, C., Wills, T. J., Burgess, N., and O’Keefe, J. (2004). Theta-modulated place-by-direction cells in the hippocampal formation in the rat. The Journal of neuroscience, 24(38):8265–8277.

Carlini, A., Bordeau, C., and Ambard, M. (2024). Auditory localization: a comprehensive practical review. Frontiers in Psychology, Volume 15-2024. doi: 10.3389/fpsyg.2024.1408073.

Chang, L. and Tsao, D. Y. (2017). The code for facial identity in the primate brain. Cell, 169(6):1013–1028.e14.

Chang, T.-Y., Doudlah, R., Kim, B., Sunkara, A., Thompson, L. W., Lowe, M. E., and Rosenberg, A. (2020). Functional links between sensory representations, choice activity, and sensorimotor associations in parietal cortex. Elife, 9:e57968.

Chillingworth, D. R. J., Marsden, J. E., and Wan, Y. H. (1982). Symmetry and bifurcation in three-dimensional elasticity, part i.Archive for rational mechanics and analysis, 80(4):295–331.

Craig, J. J. Introduction to Robotics: Mechanics and Control. Prentice Hall, third edition, (2005).

Cunningham, J. P. and Yu, B. M. (2014). Dimensionality reduction for large-scale neural recordings. Nature Neuroscience, 17 (11):1500–1509. doi: 10.1038/nn.3776.

Curto, C. and Sanderson, N. (2025). Topological neuroscience: Linking circuits to function. Annual Review of Neuroscience, 48 (Volume 48, 2025):491–518. doi:10.1146/annurev-neuro-112723-034315.

Dakin, C. J. and Rosenberg, A. Chapter3 -gravity estimation and verticality perception. In Day, B. L.and Lord, S. R., editors, Balance, Gait, and Falls, volume 159 of Handbook of Clinical Neurology, pages 43–59. Elsevier, (2018). doi:10.1016/B978-0-444-63916-5.00003-3.

DiCarlo, J. J., Zoccolan, D., and Rust, N. C. (2012). How does the brain solve visual object recognition? Neuron, 73(3):415–434.

Doudlah, R., Chang, T.-Y., Thompson, L. W., Kim, B., Sunkara, A., and Rosenberg, A. (2022). Parallel processing, hierarchical transformations, and sensorimotor associations along the ‘where’pathway. Elife, 11:e78712.

Fefferman, C., Mitter, S., and Narayanan, H. (2016). Testing the manifold hypothesis. Journal of the American Mathematical Society, 29:983–1049. doi: 10.1090/jams/852.

Finkelstein, A., Derdikman, D., Rubin, A., Foerster, J. N., Las, L., and Ulanovsky, N. (2015). Three-dimensional head-direction coding in the bat brain. Nature (London), 517(7533):159–164.

Gallego, J. A., Perich, M. G., Miller, L. E., and Solla, S. A. (2017). Neural manifolds for the control of movement. Neuron, 94(5): 978–984. doi:10.1016/j.neuron.2017.05.025.

Galletti, C., Kutz, D. F., Gamberini, M., Breveglieri, R., and Fattori, P. (2003). Role of the medial parieto-occipital cortex in the control of reaching and grasping movements. Experimental Brain Research, 153(2):158–170. doi: 10.1007/s00221-003-1589-z.

Gardner, R. J., Hermansen, E., Pachitariu, M., et al. (2022). Toroidal topology of population activity in grid cells. Nature, 602: 123–128. doi: 10.1038/s41586-021-04268-7.

Georgopoulos, A. P., Kalaska, J. F., Caminiti, R., and Massey, J. T. (1982). On the relations between the direction of two-dimensional arm movements and cell discharge in primate motor cortex. The Journal of neuroscience, 2(11):1527–1537.

Hardcastle, K., Maheswaranathan, N., Ganguli, S., and Giocomo, L. M. (2017). A multiplexed, heterogeneous, and adaptive code for navigation in medial entorhinal cortex. Neuron (Cambridge, Mass.), 94(2):375–387.e7.

Hegdé, J. and Van Essen, D. C. (2005). Role of primate visual area v4 in the processing of 3-d shape characteristics defined by disparity. Journal of Neurophysiology, 94(4):2856–2866. doi: 10.1152/jn.00802.2004.

Helmer, M., Kozyrev, V., Stephan, V., Treue, S., Geisel, T., and Battaglia, D. (2016). Model-free estimation of tuning curves and their attentional modulation, based on sparse and noisy data. PloS one, 11(1):e0146500.#x2013;.

Hinkle, D. A. and Connor, C. E. (2002). Three-dimensional orientation tuning in macaque area V4. Nat Neurosci, 5(7):665–670. doi: 10.1038/nn875.

Hirsch, M. W. Differential Topology. Springer-Verlag, (1976).

Horak, F. B. (2006). Postural orientation and equilibrium: what do we need to know about neural control of balance to prevent falls? Age and ageing, 35(suppl_2):ii7–ii11.

Hubel, D. H. and Wiesel, T. N. (1959). Receptive fields of single neurones in the cat’s striate cortex. J Physiol, 148:574–591.

Hubel, D. H. and Wiesel, T. N. (1962). Receptive fields, binocular interaction and functional architecture in the cat’s visual cortex.The Journal of Physiology, 160(1):106–154. doi: 10.1113/jphysiol.1962.sp006837.

Hubel, D. H. and Wiesel, T. N. (1968). Receptive fields and functional architecture of monkey striate cortex. J Physiol, 195(1): 215–243.

Issa, N. P., Trepel, C., and Stryker, M. P. (2000). Spatial frequency maps in cat visual cortex. Journal of Neuroscience, 20(22): 8504–8514. doi: 10.1523/JNEUROSCI.20-22-08504.2000.

Knudsen, E. I. and Konishi, M. (1978). A neural map of auditory space in the owl. Science, 200(4343):795–797. doi: 10.1126/science.644324.

Koditschek, D. E. Application of a new Lyapunov function to global adaptive attitude tracking. In Proceedings of the IEEE Conference on Decision and Control, pages 63–68, (1988).

Langdon, C., Genkin, M., and Engel, T. A. (2023). A unifying perspective on neural manifolds and circuits for cognition. Nature Reviews Neuroscience, 24(6):363–377. doi: 10.1038/s41583-023-00693-x.

Laurens, J. and Angelaki, D. E. (2019). A model-based reassessment of the three-dimensional tuning of head direction cells in rats. Journal of neurophysiology, 122(3):1274–1287.

Leigh, R. J. and Zee, D. S. The neurology of eye movements. Oxford University Press, (2015).

Lutz, O. J. and Bensmaia, S. J. (2021). Proprioceptive representations of the hand in somatosensory cortex. Current Opinion in Physiology, 21:9–16. doi:10.1016/j.cophys.2021.04.002.

Mardia, K. V. and Jupp, P. E. Directional Statistics. Wiley, (2000).

Murata, A., Gallese, V., Luppino, G., Kaseda, M., and Sakata, H. (2000). Selectivity for the shape, size, and orientation of objects for grasping in neurons of monkey parietal area aip. J Neurophysiol, 83(5):2580–2601.

O’Keefe, J. (1976). Place units in the hippocampus of the freely moving rat. Experimental Neurology, 51(1):78–109. doi:10.1016/0014-4886(76)90055-8.

Orban, G. A. (2011). The extraction of 3D shape in the visual system of human and nonhuman primates. Annual Review of Neuroscience, 34(Volume 34, 2011):361–388. doi:10.1146/annurev-neuro-061010-113819.

Page, H. J. I., Wilson, J. J., and Jeffery, K. J. (2018). A dual-axis rotation rule for updating the head direction cell reference frame during movement in three dimensions. Journal of neurophysiology, 119(1):192–208.

Rosenberg, A., Cowan, N. J., and Angelaki, D. E. (2013). The visual representation of 3D object orientation in parietal cortex.The Journal of neuroscience, 33(49):19352–19361.

Rosenberg, A., Thompson, L. W., Doudlah, R., and Chang, T.-Y. (2023). Neuronal representations supporting three-dimensional vision in nonhuman primates. Annual Review of Vision Science, 9(Volume 9, 2023):337–359. doi:10.1146/annurev-vision-111022-123857.

Rosenberg, A., Bridge, H., Connor, C. E., Samonds, J. M., Tanabe, S., and Krug, K. (2025). The richness of vision in the mammalian brain: Neural codes for visual perception and behavior in three dimensions. The Journal of Neuroscience, 45(46):e1316252025.

Secer, G., Knierim, J. J., and Cowan, N. J. (2025). Explicit error coding can mediate gain recalibration in continuous bump attractor networks. Nature communications, 16(1):9215.

Secer, G., Krishnan, B., Cowan, N. J., and Knierim, J. J. (2026). Dynamic coupling between visual landmark processing in area 29e and parahippocampal path integration circuits of the rodent cortex. bioRxiv.

Seelig, J. D. and Jayaraman, V. (2015). Neural dynamics for landmark orientation and angular path integration. Nature, 521 (7551):186–191. doi: 10.1038/nature14446.

Singh, G., Memoli, F., Ishkhanov, T., Sapiro, G., Carlsson, G., and Ringach, D. L. (2008). Topological analysis of population activity in visual cortex. Journal of vision, 8(8):11–11.

Skaggs, W., Knierim, J., Kudrimoti, H., and McNaughton, B. (1994). A model of the neural basis of the rat’s sense of direction.Advances in neural information processing systems, 7.

Stevens, K. A. (1983). Slant-tilt: The visual encoding of surface orientation. Biol Cybern, 46(3):183–195.

Swindale, N. V. (1996). Visual cortex: Looking into a Klein bottle. Curr Biol, 6(7):776–779.

Swindale, N. V. (1998). Orientation tuning curves: empirical description and estimation of parameters. Biol Cybern, 78(1):45–56.

Tanaka, S. (1995). Topological analysis of point singularities in stimulus preference maps of the primary visual cortex.Proceedings: Biological Sciences, 261(1360):pp. 81–88.

Taube, J. (1995). Head direction cells recorded in the anterior thalamic nuclei of freely moving rats. Journal of Neuroscience, 15 (1):70–86. doi: 10.1523/JNEUROSCI.15-01-00070.1995.

Taube, J., Muller, R., and Ranck, J. JB. (1990). Head-direction cells recorded from the postsubiculum in freely moving rats. I. description and quantitative analysis. The Journal of neuroscience, 10(2):420–435.

Theys, T., Pani, P., van Loon, J., Goffin, J., and Janssen, P. (2012). Selectivity for three-dimensional shape and grasping-related activity in the macaque ventral premotor cortex. Journal of Neuroscience, 32(35):12038–12050. doi: 10.1523/JNEUROSCI.1790-12.2012.

Williams, M., Street, J. S., Burgess, N., and Jeffery, K. J. (2026). Head direction cells use a head-referenced dual-axis updating rule in 3D space. Communications Biology, 9(1):1012.

Yao, M., Praturu, A., and Sharpee, T. O. (2026). Multiscale hyperbolic embedding reveals hierarchical structure in complex biological systems. NPJ systems biology and applications, 12.

Yershova, A. and LaValle, S. Deterministic sampling methods for spheres and so(3). In IEEE International Conference on Robotics and Automation, 2004. Proceedings. ICRA ‘04. 2004, volume 4, pages 3974–3980 Vol.4, (2004). doi: 10.1109/ROBOT.2004.1308891.

Yoon, I. H. R., Henselman-Petrusek, G., Yu, Y., Ghrist, R., Smith, S. L., and Giusti, C. (2024). Tracking the topology of neural manifolds across populations. Proceedings of the National Academy of Sciences, 121(46):e2407997121. doi: 10.1073/pnas.2407997121.

