## Supplementary Information for "Manifolds for the neural representation of 3D spatial experience"

### Detailed methods

***Neuronal data.*** The deductive manifold framework was applied to extracellular recordings from bats, mice, and monkeys as described below.

**3D visual orientation selectivity in macaque parietal cortex.** Neuronal recordings from the caudal intraparietal (CIP) area of rhesus monkeys (*Macaca mulatta*) were from a previous study Chang et al. (2020). In that study, data were recorded from 437 CIP neurons responding to 3D oriented planar surfaces defined by random dots with stereoscopic and perspective cues. The planes subtended 20° of visual angle, were centered on a fixation target at the center of the screen (57 cm), and were presented at a depth of 57 cm. Orientation was parameterized using tilt and slant Stevens (1983); Rosenberg et al. (2013). All combinations of tilt (0–315°, 45° steps) and slant (15–60°, 15° steps) plus the frontoparallel plane (slant = 0°, tilt undefined) were presented. Firing rates were calculated from the median visual response latency of the neurons (52 ms) to the end of the 1 s stimulus presentation.

**Head direction selectivity in the bat dorsal presubiculum.** Neuronal recordings from the dorsal presubiculum of Egyptian fruit bats was graciously provided by Arseny Finkelstein and Nachum Ulanovsky from their previous study Finkelstein et al. (2015). In that study, a custom head tracking device, consisting of a 3D arrangement of light-emitting diodes, was used to record the Euler angles of head direction of freely moving adult male bats. We used data from three experimental sessions, including two upright and one inverted. During the upright sessions, the bats crawled in a horizontal square arena ($50 \times 50\,\mathrm{cm}$) while searching for food. Inverted pitch angles were tested in between the two upright sessions by passively moving the bats into different inverted head orientations. Only cells with stable tuning in both upright sessions were analyzed.

Head direction was binned in azimuth from $-180°$ to $180°$ in $20°$ bins, and pitch was binned from $-180°$ to $180°$ in $10°$ bins. Joint azimuth–pitch firing rate maps for the upright sessions were computed by first summing the spike-count maps from the two upright sessions and summing their corresponding occupancy maps, and then dividing the summed spike-count map by the summed occupancy map. Following Finkelstein et al. (2015), bins with duration times less than 0.5 seconds were excluded. The upright and inverted firing rate maps were then concatenated.

**Head direction selectivity in the macaque anterior thalamus.** Previous work identified head direction cells in the anterior thalamus of rhesus macaques Laurens et al. (2016). To examine the cross-species generalization of our deductive framework, we included a recording from an ongoing study of this area during free exploration of an open arena. The procedures were in strict accordance with the National Institutes of Health's Guide for the Care and Use of Laboratory Animals and approved by the Institutional Animal Care and Use Committee (IACUC) at the University of Wisconsin–Madison (Protocol G005229).

*Behavioral monitoring.* The dimensions of the electrically-shielded arena were: length = 3.43 m; width = 3.43 m; height = 2.6 m. The floor, walls, door, and ceiling were matte and the walls were covered by black mesh curtains. On each wall, a DC-powered LCD screen protruded past the curtain. To encourage exploration, sucrose pellets (BioServ, Inc.) were periodically dropped from pellet dispensers in the ceiling (Lafayette, Inc.). Head movements were tracked using an eight-camera 3D motion capture system (OptiTrack, NaturalPoint, Inc.) with infrared-reflective markers affixed to the cranial implant (sampling rate = 160 Hz). The motion capture data were broadcast over a private network to the control suite (REC-GUI; RRID: SCR 019008) for real-time behavioral monitoring and saved for offline analysis Kim et al. (2019).

*Neuronal recordings.* The anterior thalamus was separated from the rest of the thalamus by the internal medullary lamina, a distinct white matter structure Jones (2007). To identify its location and plan penetration trajectories, structural magnetic resonance imaging (MRI) was performed on a 3-Tesla GE MR750 scanner. Tetrode wire bundles were positioned using a skull-mounted, wireless microdrive Baeg et al. (2021). During penetrations, white and gray matter transitions were referenced to the MRI. Neural activity was sampled at 32 kHz, digitized, and stored on a mobile neurologger (SpikeLog64; DeuteronTech, Inc.). The neuronal data were transferred and aligned with the 3D motion capture data for offline analysis. Tetrode-based spike sorting was performed using the KlustaKwik (K. Harris) semi-automatic cluster algorithm in MClust (MClust-4.0, A.D. Redish et al.) and manually refined using Offline Sorter (Plexon, Inc.).

*Head direction analysis.* To quantify head-direction tuning, head orientation was binned into 10° segments. For pitch and yaw/azimuth, bin edges ranged from –90–90° and 0–360°, respectively. Joint azimuth–pitch spike-count and duration (occupancy) maps were then calculated. If the total time at a given azimuth–pitch conjunction was less than 30 ms, that bin was excluded. The head orientation firing rate map was then calculated by dividing the spike-count map by the duration map.

**Head direction selectivity in mouse cingulum fiber bundle.** Neuronal recordings from the cingulum fiber bundle in mice were graciously provided by Dora Angelaki and Jean Laurens from a previous study Angelaki et al. (2020).

The authors recorded neuronal activity from male adult mice while they were passively positioned at different 3D orientations by a multi-axis rotator. This rotator uniformly sampled the neurons' full 3D tuning curves.

*Head direction analysis.* To avoid potential distortions associated with a particular angular parameterization, the analysis was performed directly on $\mathrm{SO}(3)$, rather than in the azimuth–tilt coordinate system defined in Angelaki et al. (2020). The space of 3D rotations was discretized into 4000 approximately uniformly distributed bins using the Super-Fibonacci sampling method from Alexa (2022), with each bin represented by a rotation matrix. At each time point, the animal's 3D head orientation was converted from the measured angular variables into a rotation matrix and assigned to the nearest $\mathrm{SO}(3)$ bin by maximizing the rotational similarity $\mathrm{tr}(R_i^\top R_j)$ between the observed rotation and each bin center. Equivalently, this selects the bin minimizing the Frobenius chord distance $\|R_i - R_j\|_F$, since $\|R_i - R_j\|_F^2 = 6 - 2\mathrm{tr}(R_i^\top R_j)$. Spike times were similarly mapped to $\mathrm{SO}(3)$ by interpolating the angular variables at each spike time, converting the resulting orientation to a rotation matrix, and assigning it to the nearest bin. Occupancy and spike-count maps were then constructed by accumulating the total time spent and the number of spikes in each $\mathrm{SO}(3)$ bin, respectively. Both maps were smoothed directly on $\mathrm{SO}(3)$ using a Gaussian kernel defined by the Frobenius chord distance between rotation matrices, and the kernel width was set to be $15°$. Finally, the 3D firing rate map was obtained by dividing the smoothed spike-count map by the smoothed occupancy map.

***Model fitting.*** In this section, we present the details for how we fit various mathematical models, which are derived in the Results section, to recorded neuronal responses.

**Fitting the Bingham function.** To fit the Bingham function on $\mathrm{S}^2$ in Eq. S35 to the responses of individual neurons, a regularization parameter was included in the optimization routine to prevent overfitting of unrealistically narrow tuning curves (large $\lambda_2$ values, see Eq. **??**). The optimization routine thus minimized the sum squared error (SSE) between the Bingham function fit ($\hat{y}_i$) and observed neuronal responses ($y_i$) plus a regularization term ($r\lambda_2^2$), where $r$ is a gain as follows:

$$\mathrm{SSE} = \sum_{i=1}^{n} (y_i - \hat{y}_i)^2 + r\lambda_2^2. \tag{S1}$$

The optimal value of $r$ was found for each neuron using k-fold cross-validation. The optimization was performed using *fmincon* in MATLAB.

**Fitting the CMWK function.** To fit the CMWK function on $\mathrm{T}^2$ in Eq. **??** to the firing rate maps of individual head direction cells, nonlinear least-squares optimization was performed using *lsqnonlin* in MATLAB to minimize the SSE between the function fit and observed responses. In the fitting for 2 DOF head direction, we multiplied the CMWK exponential term with the constant $\exp(-\mathrm{Tr}[P])$, i.e.:

$$\tilde{\varphi}(R) = dc + G \cdot \exp(\mathrm{Tr}[PR]) \cdot \exp(-\mathrm{Tr}[P]), \tag{S2}$$

so that $G$ approximates the maximum firing rate.

***Data informed model refinement.*** These manifold-compatible tuning functions derived using a deductive approach *fully* parameterize responses over the manifold, but this parameterization may not be optimal for observed neuronal activity. Therefore, we next assess whether the proposed parameterization captures the features present in the data (Fig. **??**B). As an example, we test whether the full Bingham function for planar surfaces is necessary to model 3D orientation responses in area CIP.

Previously published findings in area CIP consistently report that the majority of Bingham function fits result in the $\rho$ parameter being near $90°$, indicating that the tuning anisotropy is approximately aligned with the slant–tilt axis Rosenberg et al. (2013); Chang et al. (2020); Doudlah et al. (2022). Therefore, we wanted to assess if neuronal responses in area CIP were better described by the Bingham function with $\rho$ as a free parameter (full model) or fixed ($\rho = 90°$; reduced model).

To test this, we fit the 3D orientation tuning curves of 437 CIP neurons Chang et al. (2020); Doudlah et al. (2022) using the full and reduced Bingham functions (see Detailed methods). To directly compare the performance of each model, Akaike's Information Criterion (AIC) corrected for finite sample size was calculated Burnham and Anderson (2002). Across the population, we found that $37\%$ of neuronal responses were better described by the full model. To compare neuronal tuning when the full or reduced model was a better fit, we compared the joint distribution of $\rho$ and $\lambda_1$ (anisotropy) parameters (Fig. S3B). As expected, when the fit $\rho$ value from the full model was close to $90°$, the reduced model was a better fit. Furthermore, fits with $\lambda_1 \approx 0$ (i.e., isotropic tuning) were also better described by the reduced model since $\rho$ becomes undefined as $\lambda_1$ approaches zero. We found that the reduced model better described neuronal responses when the true $\rho$ term, from the full model, was within $\pm 25°$ (Fig. S3C).

Given that a large proportion of neurons require the $\rho$ parameter to accurately model 3D orientation responses in area CIP, we continue to use the full Bingham function. This is consistent with a previous report that found that a combination of $\lambda_1$ and $\rho$ were jointly required to appropriately model CIP responses (the full model was a better fit for $88\%$ of the neurons) Rosenberg et al. (2013). We now consider how the submanifold changes when orientation is restricted within the frontoparallel plane, and show that a commonly used model for describing orientation in the frontoparallel plane can be derived from the Bingham function on this submanifold.

***Comparing the manifold structure for 3D surface orientation.*** We used principal components analysis (PCA) to infer the manifold geometry of 3D visual planar surface orientation from CIP pseudopopulation responses. First, average z-scored firing rates for each orientation (N = 33) and neuron (N = 437) were computed at each of the four stimulus distances. Second, PCA was performed on the resulting response matrices (33 x 437). The first three principal components (PCs; variance explained: 37 cm = 0.57; 57 cm = 0.64; 97 cm = 0.52; 137 cm = 0.43) were used for subsequent analyses.

To directly compare the geometry of the inductive manifold on which those PCs resided to the deductive (antipodally symmetric, spherical) manifold derived in the Results, we fit an affine transformation that best mapped the first three PCs to their corresponding points (i.e., slant–tilt combinations) on the deductive manifold:

$$\begin{bmatrix}\hat{x_i}\\ \hat{y_i}\\ \hat{z_i}\end{bmatrix} = \begin{bmatrix}a_{1,1} & a_{1,2} & a_{1,3}\\ a_{2,1} & a_{2,2} & a_{2,3}\\ a_{3,1} & a_{3,2} & a_{3,3}\end{bmatrix}\begin{bmatrix}x_i\\ y_i\\ z_i\end{bmatrix} + \begin{bmatrix}t_x\\ t_y\\ t_z\end{bmatrix} \tag{S3}$$

where elements $a_{i,j}$ comprise a general 3×3 transformation matrix, and $t_{x,y,z}$ is a three-dimensional translation vector. The parameters were fit to minimize the sum squared Euclidean distance between points $(\hat{x_i}, \hat{y_i}, \hat{z_i})$ and their locations $(x_i', y_i', z_i')$ on an antipodally symmetric sphere, where $i$ indicates a unique orientation. The optimization was performed by calculating the least-squares solution in MATLAB. The fitted matrix was nonsingular, so the affine transformation defines a smooth, invertible local change of coordinates.

### 3D rotation representations and conversions

Here, we introduce the representations used to parameterize rotations in 3D space, including Euler angles, quaternions, and rotation matrices. We also present the conversions between these representations, which serve as the foundation for the mathematical derivations that follow.

***Euler angle parameterization of rotation matrices in 3D space.*** Here are the $3 \times 3$ rotation matrices about the $x$, $y$, and $z$ axis in $\mathbb{R}^3$:

$$R_x(\phi) = \begin{bmatrix}1 & 0 & 0\\ 0 & \cos\phi & -\sin\phi\\ 0 & \sin\phi & \cos\phi\end{bmatrix},$$

$$R_y(\theta) = \begin{bmatrix}\cos\theta & 0 & \sin\theta\\ 0 & 1 & 0\\ -\sin\theta & 0 & \cos\theta\end{bmatrix}, \tag{S4}$$

$$R_z(\psi) = \begin{bmatrix}\cos\psi & -\sin\psi & 0\\ \sin\psi & \cos\psi & 0\\ 0 & 0 & 1\end{bmatrix},$$

where $\phi$, $\theta$, $\psi$ are referred to as roll, pitch, yaw, respectively. For the rotation matrices describing slant ($s$), a rotation about the $y$-axis, and tilt ($t$), a rotation about the $z$-axis, the variables $s$ and $t$ can be substituted in for $\theta$ and $\psi$, respectively.

The $3 \times 3$ rotation matrix for rotations of roll, pitch, and yaw is described by

$$\begin{aligned} R^{3\times3} &= R_z(\psi)R_y(\theta)R_x(\phi)\\ &= \begin{bmatrix}\cos\theta\cos\psi & \sin\phi\sin\theta\cos\psi - \cos\phi\sin\psi & \cos\phi\sin\theta\cos\psi + \sin\phi\sin\psi\\ \cos\theta\sin\psi & \cos\phi\cos\psi + \sin\phi\sin\theta\sin\psi & \cos\phi\sin\theta\sin\psi - \sin\phi\cos\psi\\ -\sin\theta & \sin\phi\cos\theta & \cos\phi\cos\theta\end{bmatrix}. \end{aligned} \tag{S5}$$

In the derivation of the Bingham function on $\mathrm{SO}(3)$, parameters $\alpha, \beta, \gamma$ are substituted for $\phi, \theta, \psi$, respectively.

***Converting Euler angles to quaternions.*** To convert Euler angles, roll ($\phi$), pitch ($\theta$), and yaw ($\psi$), to quaternions, assuming the standard Hamilton convention, where $\mathbf{q} = w + xi + yj + zk$, the quaternion can be written as

$$\mathbf{q} = \begin{bmatrix} w \\ x \\ y \\ z \end{bmatrix} = \begin{bmatrix} \cos(\phi/2)\cos(\theta/2)\cos(\psi/2) + \sin(\phi/2)\sin(\theta/2)\sin(\psi/2) \\ \sin(\phi/2)\cos(\theta/2)\cos(\psi/2) - \cos(\phi/2)\sin(\theta/2)\sin(\psi/2) \\ \cos(\phi/2)\sin(\theta/2)\cos(\psi/2) + \sin(\phi/2)\cos(\theta/2)\sin(\psi/2) \\ \cos(\phi/2)\cos(\theta/2)\sin(\psi/2) - \sin(\phi/2)\sin(\theta/2)\cos(\psi/2) \end{bmatrix}. \tag{S6}$$

***Converting quaternions to rotation matrices.*** To convert a unit quaternion ($\mathbf{q}$) into a rotation matrix, assuming the standard Hamilton convention, the rotation matrix ($R_q$) can be written as

$$R_q = \begin{bmatrix} 1-2y^2-2z^2 & 2xy-2zw & 2xz+2yw \\ 2xy+2zw & 1-2x^2-2z^2 & 2yz-2xw \\ 2xz-2yw & 2yz+2xw & 1-2x^2-2y^2 \end{bmatrix}. \tag{S7}$$

### Mathematical derivations

***Bingham function on $\mathrm{SO}(3)$: $\mu$ vectors.*** As described in the main text, the $\mu$ vectors (i.e., columns of $M$) are given by $M = M^*\bar{R}(\alpha,\beta,\gamma)J$, where $\bar{R}$ is given by Eq. **??** and Eq. S5, and $M^*$, as defined in Eq. **??**, is given by

$$M^* = \begin{bmatrix} w^* & -x^* & -y^* & -z^* \\ x^* & w^* & -z^* & y^* \\ y^* & z^* & w^* & -x^* \\ z^* & -y^* & x^* & w^* \end{bmatrix}. \tag{S8}$$

In $M^*$, $w^*, x^*, y^*, z^*$ are defined by Eq. S6 for a preferred roll ($\phi^*$), pitch ($\theta^*$), and yaw ($\psi^*$). The columns of $M^*\bar{R}(\alpha,\beta,\gamma)$, in order left-to-right, are as follows:

$$\begin{aligned}
\mu_4 &= \begin{bmatrix} w^* \\ x^* \\ y^* \\ z^* \end{bmatrix}, \\
\mu_1 &= \begin{bmatrix} -x^*\cos\beta\cos\gamma - y^*\cos\beta\sin\gamma + z^*\sin\beta \\ w^*\cos\beta\cos\gamma - z^*\cos\beta\sin\gamma - y^*\sin\beta \\ z^*\cos\beta\cos\gamma + w^*\cos\beta\sin\gamma + x^*\sin\beta \\ -y^*\cos\beta\cos\gamma + x^*\cos\beta\sin\gamma - w^*\sin\beta \end{bmatrix}, \\
\mu_2 &= \begin{bmatrix} -x^*\left(\sin\alpha\sin\beta\cos\gamma - \cos\alpha\sin\gamma\right) - y^*\left(\cos\alpha\cos\gamma + \sin\alpha\sin\beta\sin\gamma\right) - z^*\sin\alpha\cos\beta \\ w^*\left(\sin\alpha\sin\beta\cos\gamma - \cos\alpha\sin\gamma\right) - z^*\left(\cos\alpha\cos\gamma + \sin\alpha\sin\beta\sin\gamma\right) + y^*\sin\alpha\cos\beta \\ z^*\left(\sin\alpha\sin\beta\cos\gamma - \cos\alpha\sin\gamma\right) + w^*\left(\cos\alpha\cos\gamma + \sin\alpha\sin\beta\sin\gamma\right) - x^*\sin\alpha\cos\beta \\ -y^*\left(\sin\alpha\sin\beta\cos\gamma - \cos\alpha\sin\gamma\right) + x^*\left(\cos\alpha\cos\gamma + \sin\alpha\sin\beta\sin\gamma\right) + w^*\sin\alpha\cos\beta \end{bmatrix}, \\
\mu_3 &= \begin{bmatrix} -x^*\left(\cos\alpha\sin\beta\cos\gamma + \sin\alpha\sin\gamma\right) - y^*\left(\cos\alpha\sin\beta\sin\gamma - \sin\alpha\cos\gamma\right) - z^*\cos\alpha\cos\beta \\ w^*\left(\cos\alpha\sin\beta\cos\gamma + \sin\alpha\sin\gamma\right) - z^*\left(\cos\alpha\sin\beta\sin\gamma - \sin\alpha\cos\gamma\right) + y^*\cos\alpha\cos\beta \\ z^*\left(\cos\alpha\sin\beta\cos\gamma + \sin\alpha\sin\gamma\right) + w^*\left(\cos\alpha\sin\beta\sin\gamma - \sin\alpha\cos\gamma\right) - x^*\cos\alpha\cos\beta \\ -y^*\left(\cos\alpha\sin\beta\cos\gamma + \sin\alpha\sin\gamma\right) + x^*\left(\cos\alpha\sin\beta\sin\gamma - \sin\alpha\cos\gamma\right) + w^*\cos\alpha\cos\beta \end{bmatrix}.
\end{aligned} \tag{S9}$$

***Tuning Shape Analysis of the CMWK function.*** Here we derive the parameters that define the tuning shape of the CMWK function. In $\mathrm{SO}(3)$, a small rotation away from the current orientation $R_0$ takes the form

$$R(t) = R_0 \exp(t\hat{\omega}), \tag{S10}$$

where $t$ is the rotation time, $||\omega||$ is the rotation velocity, $\frac{\omega}{||\omega||}$ is the rotation axis, and $\hat{\omega}$ is the skew-symmetric matrix with $\hat{\omega} \cdot v = \omega \times v$. The exponential map is given by the Rodrigues formula.

We will use two identities derived from direct calculation: for any $A \in \mathbb{R}^{3\times3}$ and $\omega \in \mathbb{R}^3$,

$$\mathrm{Tr}[A\hat{\omega}] = (A^\top - A)^\vee \cdot \omega, \qquad \hat{\omega}^2 = \omega\omega^\top - \|\omega\|^2 I, \tag{S11}$$

where $^\vee$ extracts the associated $3 \times 1$ vector from a skew-symmetric matrix.

**Gradient and critical points.** Consider the CMWK function $\varphi(R) = \mathrm{Tr}[PR]$ with $P$ symmetric. Along the one-parameter curve $R(t)$ as defined in Eq. S10, the first derivative is

$$\begin{aligned}\frac{d}{dt}\varphi(R(t)) &= \frac{d}{dt}\mathrm{Tr}\left[PR_0\exp(t\hat{\omega})\right] \\ &= \mathrm{Tr}\left[PR_0\frac{d}{dt}\exp(t\hat{\omega})\right].\end{aligned} \tag{S12}$$

Since $\frac{d}{dt}\exp(t\hat{\omega})\Big|_{t=0} = \hat{\omega}$, we have

$$\begin{aligned}\frac{d\varphi}{dt}\Big|_{t=0} &= \mathrm{Tr}[PR_0\hat{\omega}] \\ &= (R_0^\top P - PR_0)^\vee \cdot \omega.\end{aligned} \tag{S13}$$

Thus, the gradient in this local $\omega$-coordinates is $\nabla\varphi(R_0) = (R_0^\top P - PR_0)^\vee$.

When $R_0$ is a critical point of $\varphi$, $\nabla\varphi = 0$, thus $R_0^\top P = PR_0$. Since $P$ is symmetric, $R_0^\top P = (PR_0)^\top$, so $(PR_0)^\top = PR_0$. Therefore, $R_0$ is a critical point of $\varphi$ if and only if $PR_0$ is symmetric, agreeing with Koditschek (1988).

**Hessian at critical points.** The second derivative along the curve $R(t)$ is:

$$\frac{d^2}{dt^2}\varphi(R(t))\Big|_{t=0} = \frac{d^2}{dt^2}\mathrm{Tr}[PR_0\exp(t\hat{\omega})]\Big|_{t=0} \tag{S14}$$

Using the Taylor expansion $\exp(t\hat{\omega}) = I + t\hat{\omega} + \frac{t^2}{2}\hat{\omega}^2 + O(t^3)$:

$$\begin{aligned}\varphi(R(t)) &= \mathrm{Tr}\left[PR_0\left(I + t\hat{\omega} + \frac{t^2}{2}\hat{\omega}^2 + O(t^3)\right)\right] \\ &= \mathrm{Tr}[PR_0] + t\,\mathrm{Tr}[PR_0\hat{\omega}] + \frac{t^2}{2}\mathrm{Tr}[PR_0\hat{\omega}^2] + O(t^3)\end{aligned} \tag{S15}$$

Therefore,

$$\begin{aligned}\frac{d^2\varphi}{dt^2}\Big|_{t=0} &= \mathrm{Tr}[PR_0\hat{\omega}^2] \\ &= \mathrm{Tr}[PR_0(\omega\omega^\top - \|\omega\|^2 I)] \\ &= \omega^\top PR_0\,\omega - \mathrm{Tr}[PR_0]\|\omega\|^2 \\ &= \omega^\top(PR_0 - \mathrm{Tr}[PR_0]\cdot I)\omega.\end{aligned} \tag{S16}$$

Thus, the Hessian at critical points in $\omega$-coordinates is $H = PR_0 - \mathrm{Tr}[PR_0]\cdot I$. Furthermore, if $PR_0$ has eigenvalues $\lambda_1, \lambda_2, \lambda_3$, we have $H = PR_0 - (\lambda_1+\lambda_2+\lambda_3)I$. Let $v_i$ be an eigenvector of $PR_0$ corresponding to eigenvalue $\lambda_i$, i.e., $PR_0 v_i = \lambda_i v_i$, where $i = 1,2,3$. Multipling $H$ with $v_i$, we have

$$\begin{aligned}Hv_i &= (PR_0 - (\lambda_1+\lambda_2+\lambda_3)I)\,v_i \\ &= PR_0 v_i - (\lambda_1+\lambda_2+\lambda_3)v_i \\ &= \lambda_i v_i - (\lambda_1+\lambda_2+\lambda_3)v_i \\ &= (\lambda_i - (\lambda_1+\lambda_2+\lambda_3))\,v_i.\end{aligned} \tag{S17}$$

So $v_i$ is also an eigenvector of $H$, with eigenvalue $h_i = \lambda_i - (\lambda_1+\lambda_2+\lambda_3)$. Therefore, the eigenvalues of $H$ are:

$$\begin{aligned}h_1 &= \lambda_1 - (\lambda_1+\lambda_2+\lambda_3) = -\lambda_2 - \lambda_3 \\ h_2 &= \lambda_2 - (\lambda_1+\lambda_2+\lambda_3) = -\lambda_1 - \lambda_3 \\ h_3 &= \lambda_3 - (\lambda_1+\lambda_2+\lambda_3) = -\lambda_1 - \lambda_2\end{aligned} \tag{S18}$$

This implies that $R_0$ is a maximum of the CMWK function only if all the eigenvalues $h_1$, $h_2$ and $h_3$ are nonnegative (strictly positive for uniqueness).

**Critical points and their classification.** Denote the distinct eigenvalues of $P$ by $\pi_1, \pi_2, \pi_3$, with $\pi_1 > \pi_2 > \pi_3$ and $\pi_2 + \pi_3 > 0$, and the corresponding eigenvectors by $v_1$, $v_2$ and $v_3$. Then eigenvalue decomposition gives $P = V\,\mathrm{diag}(\pi_1, \pi_2, \pi_3)\,V^\top$, where $V = [v_1\; v_2\; v_3]$ is orthonormal. Because $P$ has distinct eigenvalues, the condition that

$PR_0$ is symmetric implies that $R_0$ must take the form $V\,\mathrm{diag}(\pm1,\pm1,\pm1)\,V^\top$ with $\det=+1$, giving four critical points, as summarized in Table 1.

**Table 1.** Critical points of the CMWK function and their classification.

| Critical point $R_0$ | $\varphi(R_0)$ | Hessian eigenvalues $h_1$, $h_2$, $h_3$ | Type |
|---|---|---|---|
| $V\,\mathrm{diag}(1,1,1)\,V^\top$ | $\pi_1+\pi_2+\pi_3$ | $-\pi_2-\pi_3,\ -\pi_1-\pi_3,\ -\pi_1-\pi_2$ | max |
| $V\,\mathrm{diag}(1,-1,-1)\,V^\top$ | $\pi_1-\pi_2-\pi_3$ | $\pi_2+\pi_3,\ \pi_3-\pi_1,\ \pi_2-\pi_1$ | saddle |
| $V\,\mathrm{diag}(-1,1,-1)\,V^\top$ | $-\pi_1+\pi_2-\pi_3$ | $\pi_3-\pi_2,\ \pi_1+\pi_3,\ \pi_1-\pi_2$ | saddle |
| $V\,\mathrm{diag}(-1,-1,1)\,V^\top$ | $-\pi_1-\pi_2+\pi_3$ | $\pi_2-\pi_3,\ \pi_1-\pi_3,\ \pi_1+\pi_2$ | min |

**Tuning sharpness at the maximum (preferred head orientation).** At the maximum $R_0=I$, the Taylor expansion of $\varphi$ along a $\xi$-parameterized curve is

$$\varphi(R)\approx\varphi_{\max}+\nabla\varphi(R_0)^\top\xi+\frac{1}{2}\xi^\top H\xi,\qquad R=R_0\exp(\hat{\xi}). \tag{S19}$$

Since $R_0=I$ is a critical point, $\nabla\varphi(R_0)=0$, we are left with

$$\varphi(R)\approx\varphi_{\max}+\frac{1}{2}\xi^\top H\xi, \tag{S20}$$

where $\varphi_{\max}=\pi_1+\pi_2+\pi_3$ and $H=P-\mathrm{Tr}[P]\cdot I$ from Eq. S16. Let $\xi=\xi_1v_1+\xi_2v_2+\xi_3v_3$. Since $H$ is diagonal in the eigenbasis of $P$, we have $\xi^\top H\xi=h_1\xi_1^2+h_2\xi_2^2+h_3\xi_3^2$. Substituting into Eq. S20, we have:

$$\varphi(R)\approx\varphi_{\max}+\frac{1}{2}\big(h_1\xi_1^2+h_2\xi_2^2+h_3\xi_3^2\big). \tag{S21}$$

For a rotation purely about $v_i$ by angle $\theta$, we have $\xi=\theta v_i$, so

$$\begin{aligned}\varphi(R)&\approx\varphi_{\max}+\frac{1}{2}h_i\theta^2\\&=\varphi_{\max}-\frac{1}{2}|h_i|\theta^2.\end{aligned} \tag{S22}$$

This is a downward parabola in $\theta$, and $|h_i|$ controls how fast $\varphi$ drops as the head rotates away from the preferred orientation. Larger $|h_i|$ means a steeper drop, i.e. sharper tuning.
Since $\pi_1>\pi_2>\pi_3$, we have $|h_1|<|h_2|<|h_3|$. Therefore, the axis with the largest eigenvalue of $P$ corresponds to the broadest tuning, and vice versa. That is:

- Rotating about $v_1$ (largest $\pi_1$): broadest tuning, sharpness $=\pi_2+\pi_3$.
- Rotating about $v_3$ (smallest $\pi_3$): sharpest tuning, sharpness $=\pi_1+\pi_2$.
- The anisotropy ratio is $(\pi_1+\pi_2)/(\pi_2+\pi_3)$.

***Equivalence of the Bingham and CMWK function.*** In this section, we show that the Bingham and CMWK distributions are equivalent on $\mathrm{S}^3$. We begin by decomposing the Bingham DOFs into three parameters for the preferred orientation and six for the tuning shape, matching the structure of the CMWK distribution. Then we derive the conversion from CMWK parameters to Bingham parameters.

**Decomposition of the Bingham function into local error coordinates.** In a “global coordinate" frame, the $A$ matrix in the quadratic term of the Bingham function, $\mathbf{x}^T A\mathbf{x}$, has 10 independent components, and $\mathbf{x}$ is a quaternion $\mathbf{q}$. However, due to the unit constraint $\mathbf{q}^T\mathbf{q}=1$,

$$\mathbf{q}^\top(A+\lambda I_4)\mathbf{q}=\mathbf{q}^\top A\mathbf{q}+\lambda, \tag{S23}$$

giving

$$G\exp\Big(\mathbf{q}^\top(A+\lambda I_4)\mathbf{q}\Big)=\Big(Ge^{\lambda}\Big)\exp\Big(\mathbf{q}^\top A\mathbf{q}\Big). \tag{S24}$$

Hence, adding $\lambda I_4$ to $A$ simply rescales the gain $G$, and does not change the shape of the function on $\mathrm{S}^3$. For simplicity (without loss of generality), we keep the maximum eigenvalue of $A$ to be $0$, i.e. $\lambda_4=0$, as mentioned in the main text, thus leaving $A$ with 9 DOFs.

To transform the representation into a “local coordinate" frame, we let $\mathbf{q}^*$, a 4-vector or quaternion, be the preferred orientation and $\mathbf{q}$ be the current orientation. We define the delta quaternion $\Delta\mathbf{q}$ as the orientation error, $\Delta\mathbf{q} = {\mathbf{q}^*}^{-1} \otimes \mathbf{q}$, where $\otimes$ is the quaternion multiplication, i.e., for two quaternions $\mathbf{p} = (w_p, p)$ and $\mathbf{q} = (w_q, q)$ with $p = (x_p, y_p, z_p)$ and $q = (x_q, y_q, z_q)$, $p \otimes q = (w_p w_q - p \cdot q,\ w_p q + w_q p + p \times q)$. Using the matrix representation of quaternion left-multiplication, we can write $\mathbf{q} = Q^* \Delta\mathbf{q}$, where $Q^*$ is a $4 \times 4$ orthogonal matrix constructed from the components of $\mathbf{q}^*$, same as Eq. **??**. Substituting the expression for $\mathbf{q}$ into the global quadratic form:

$$\begin{aligned} \mathbf{q}^\top A \mathbf{q} &= (Q^* \Delta\mathbf{q})^\top A (Q^* \Delta\mathbf{q}) \\ &= \Delta\mathbf{q}^\top ({Q^*}^\top A Q^*) \Delta\mathbf{q} \\ &= \Delta\mathbf{q}^\top A_{\text{local}} \Delta\mathbf{q} \end{aligned} \tag{S25}$$

where $A_{\text{local}} = {Q^*}^\top A Q^*$ is the representation of $A$ in the coordinate system centered at the identity quaternion $\mathbf{q}_I = [1, 0, 0, 0]^\top$.

The decomposition reveals a clear split in the 9 total DOFs. First, the preferred orientation (3-DOF) is captured by the choice of $\mathbf{q}^*$, which defines $Q^*$. Second, the shape of the tuning curve at the preferred orientation is described by $A_{\text{local}}$. Since the peak is fixed at the identity, the first eigenvector of $A_{\text{local}}$ is constrained to $[1, 0, 0, 0]^T$. This forces the first row and column of $A_{\text{local}}$ to be zero (excluding the scalar offset as the first element), leaving a $3 \times 3$ symmetric sub-matrix that describes the shape and orientation of the “ellipsoidal tuning" in the local space, giving the remaining 6 DOFs.

To see the second point mathematically, first note that when $\mathbf{q} = \mathbf{q}^*$, $\mathbf{q}^\top A \mathbf{q}$ reaches its maximum. Therefore, according to the Rayleigh-Ritz Theorem, $\mathbf{q}^*$ is the eigenvector of $A$ that corresponds to the maximum eigenvalue, denoted by $\lambda_{max}$. Since $\mathbf{q}$ and $\mathbf{q}^*$ are unit quaternions, $Q^*$ is orthogonal, that is, ${Q^*}^{-1} = {Q^*}^\top$. Hence $A = Q^* A_{\text{local}} {Q^*}^\top$, and thus

$$A\mathbf{q}^* = Q^* A_{\text{local}} {Q^*}^\top \mathbf{q}^*. \tag{S26}$$

Because $Q^*$ rotates the identity $\mathbf{q}_I$ to $\mathbf{q}^*$, we have $Q^* \mathbf{q}_I = \mathbf{q}^*$. Therefore,

$$\begin{aligned} A\mathbf{q}^* &= Q^* A_{\text{local}} \mathbf{q}_I \\ &= Q^* a_{\text{local}}, \end{aligned} \tag{S27}$$

where $a_{\text{local}}$ is the first column of $A_{\text{local}}$. Thus, we have

$$\begin{aligned} Q^* a_{\text{local}} &= \lambda_{max} \mathbf{q}^* \\ &= \lambda_{max} Q^* \mathbf{q}_I \\ &= Q^* (\lambda_{max} \mathbf{q}_I). \end{aligned} \tag{S28}$$

Since $Q^*$ is orthogonal, it must be full rank, indicating that $a_{\text{local}} = \lambda_{max} \mathbf{q}_I$. As a result, $A_{\text{local}}$ is a block diagonal matrix with the maximum eigenvalue of $A$ at the top-left corner.

Therefore, by expressing the tuning function in terms of $\Delta q$, the preferred orientation is “pulled out" of the matrix $A$, and the Bingham function on $\mathrm{S}^3$ is rewritten in a similar form to the CMWK function on $\mathrm{SO}(3)$—both $A_{\text{local}}$ and $P$ have 6 DOFs, while $\delta q$ and $R$ contain the remaining 3 DOFs for the preferred orientation.

**Conversion from CMWK Parameters to Bingham Parameters.** Here, we show how the parameters of the CMWK model can be converted into the parameters of a Bingham function on the unit quaternion sphere $\mathrm{S}^3$. Recall the CMWK function is defined as in Eq. **??**, with $P = P^\top \in \mathbb{R}^{3\times3}$, $R \in \mathrm{SO}(3)$, and $dc, G \in \mathbb{R}$. The corresponding Bingham function is written as

$$\Psi(\mathbf{q}) = dc + G \exp\left(\mathbf{q}^\top A \mathbf{q}\right), \tag{S29}$$

where $\mathbf{q} \in \mathbb{R}^4$ is a unit quaternion and $A = A^\top \in \mathbb{R}^{4\times4}$.

$P$ has the form

$$P = \begin{bmatrix} p_{11} & p_{12} & p_{13} \\ p_{12} & p_{22} & p_{23} \\ p_{13} & p_{23} & p_{33} \end{bmatrix}. \tag{S30}$$

Let $\mathbf{q}^* \in \mathrm{S}^3$ be the quaternion corresponding to the preferred orientation and let $\mathbf{q} \in \mathrm{S}^3$ be the current orientation. We define the local orientation error quaternion as $\Delta\mathbf{q} = {\mathbf{q}^*}^{-1} \otimes \mathbf{q}$. Recall from the last section, left quaternion multiplication gives $\mathbf{q} = Q^* \Delta\mathbf{q}$, with $Q^* \in \mathbb{R}^{4\times4}$. Setting $R$ as a function of $\Delta\mathbf{q}$ as in Eq. S7, the exponent in Eq. **??**

can be written as a quadratic form in $\Delta\mathbf{q}$:

$$\mathrm{Tr}(PR(\Delta\mathbf{q})) = \Delta\mathbf{q}^\top \mathcal{K}(P)\Delta\mathbf{q}, \tag{S31}$$

where the linear map $\mathcal{K} : \mathbb{R}^{3\times3}_{\mathrm{sym}} \to \mathbb{R}^{4\times4}_{\mathrm{sym}}$ is

$$\mathcal{K}(P) = \begin{bmatrix} \mathrm{Tr}(P) & 0 & 0 & 0 \\ 0 & p_{11}-p_{22}-p_{33} & 2p_{12} & 2p_{13} \\ 0 & 2p_{12} & -p_{11}+p_{22}-p_{33} & 2p_{23} \\ 0 & 2p_{13} & 2p_{23} & -p_{11}-p_{22}+p_{33} \end{bmatrix}. \tag{S32}$$

Thus, in the local coordinate frame centered at the identity quaternion $\mathbf{q}_I = [1,0,0,0]^\top$, the Bingham matrix is $A_{\mathrm{local}} = \mathcal{K}(P)$.

Since $A_{\mathrm{local}} = \mathcal{K}(P)$, from Eq. S25, the conversion from CMWK parameters, denoted by $(P,\psi^*,\theta^*,\phi^*,dc_{\mathrm{CMWK}},G_{\mathrm{CMWK}})$, to Bingham parameters $(A,dc_{\mathrm{Bingham}},G_{\mathrm{Bingham}})$ is

$$A = Q^*\mathcal{K}(P)Q^{*\top}, \tag{S33}$$

$$dc_{\mathrm{Bingham}} = dc_{\mathrm{CMWK}}, \qquad G_{\mathrm{Bingham}} = G_{\mathrm{CMWK}}. \tag{S34}$$

***Bingham function on lower dimensions.***

**Bingham function on $\mathrm{S}^2$ for an achiral bar.** The derivation of the Bingham function on $\mathrm{S}^2$ follows the procedure described for $\mathrm{SO}(3)$. Briefly, the function $\Psi : \mathrm{S}^2 \to \mathbb{R}^1$ maps points from the 2-sphere to the real number line, and has the form $\Psi(\mathbf{x}) = \exp(\mathbf{x}^\top A\mathbf{x})$. Here, $\mathbf{x}$ is a 3-vector describing points on the unit sphere and $A$ is a $3\times3$ symmetric matrix. Simplifying the expression:

$$\Psi(\mathbf{x}) = dc + G\cdot\exp\left(\lambda_1(\mu_1^\top\mathbf{x})^2 + \lambda_2(\mu_3^\top\mathbf{x})^2\right), \tag{S35}$$

where $\mu_{1,3}$ are columns of the matrix of eigenvectors of $A$. Denote the eigenvalues of $A$ by $\xi_1$, $\xi_2$ and $\xi_3$, where $\xi_3 \geq \xi_2 \geq \xi_1$, then $\lambda_1 = \xi_1 - \xi_2$ and $\lambda_2 = \xi_3 - \xi_2$ are differences between them. Here, $\lambda_2$ set the bandwidth and $\lambda_1$ sets the aspect ratio of the tuning curve Rosenberg et al. (2013).

The Bingham function is formed by taking the intersection of an ellipsoid with a unit sphere, and $\mu_{1,2,3}$ aligns with the principal axes of the ellipsoid. According to Rayleigh–Ritz theorem, the Bingham function reaches its maximum at $x = \mu_3$, so $\mu_3$ corresponds to the preferred orientation determined by the preferred slant and tilt $(s^*,t^*)$:

$$\mu_3 = \begin{bmatrix} \cos s^*\cos t^* \\ \cos s^*\sin t^* \\ -\sin s^* \end{bmatrix}. \tag{S36}$$

The remaining vectors $\mu_1$ and $\mu_2$ span the plane orthogonal to $\mu_3$ and determine the principal directions of falloff away from the preferred orientation. Their orientation within this plane is set by an additional angle $\rho$, corresponding to a rotation about $\mu_3$.

To construct the full orthonormal frame, we begin with a reference bar aligned with the $x$-axis. First, we rotate the other two axes about the reference bar axis by $R_x(\rho)$. This fixes the falloff directions of the firing rate relative to the bar. We then apply the slant–tilt rotation $R_{s,t}(s^*,t^*)$, which maps the reference $x$-axis to the preferred direction $\mu_3$. Thus, the frame for the bar is given by

$$M_B = R_{s,t}(s^*,t^*)R_x(\rho)I^{3\times3}. \tag{S37}$$

In this construction, the columns of $M_B$ are initially ordered as $\mu_3,\mu_1,\mu_2$. Since $M_B$ is orthonormal, we can reorder the columns and write $M_B = [\mu_1, \mu_2, \mu_3]$. This gives

$$\mu_1 = \begin{bmatrix} \sin\rho\sin s^*\cos t^* - \cos\rho\sin t^* \\ \sin\rho\sin s^*\sin t^* + \cos\rho\cos t^* \\ \sin\rho\cos s^* \end{bmatrix},$$
$$\mu_2 = \begin{bmatrix} \cos\rho\sin s^*\cos t^* + \sin\rho\sin t^* \\ \cos\rho\sin s^*\sin t^* - \sin\rho\cos t^* \\ \cos\rho\cos s^* \end{bmatrix}, \tag{S38}$$
$$\mu_3 = \begin{bmatrix} \cos s^*\cos t^* \\ \cos s^*\sin t^* \\ -\sin s^* \end{bmatrix}.$$

**Bingham function on $\mathrm{S}^1$.** For an achiral bar on $\mathrm{S}^1$, the rotation matrix describing its planar orientation is

$$R_z(\psi) = \begin{bmatrix} \cos\psi & -\sin\psi \\ \sin\psi & \cos\psi \end{bmatrix}. \tag{S39}$$

Starting with a function $\Psi : \mathrm{S}^1 \to \mathbb{R}^1$, the Bingham function on $\mathrm{S}^1$ can be written as

$$\Psi(\mathbf{x}) = dc + G \cdot \exp\left(\lambda(\mu^\top \mathbf{x})^2\right). \tag{S40}$$

Here, $\mathbf{x} = [\cos\psi, \sin\psi]^\top$ is a unit vector on $\mathrm{S}^1$, $\mu = [\cos\psi^*, \sin\psi^*]^\top$ is the first column of $R_z(\psi)$ (Eq. S39), where $\psi^*$ defines the preferred orientation, and $\lambda$ is the bandwidth. $G$ is a gain, and $dc$ is an offset. Substituting $\mathbf{x}$ into Eq. S40 gives

$$\Psi(\psi) = dc + G \cdot \exp\left(\lambda\cos^2(\psi - \psi^*)\right). \tag{S41}$$

Using $\cos^2(\psi - \psi^*) = \frac{1+\cos(2(\psi-\psi^*))}{2}$, this can be rewritten as

$$\Psi(\psi) = dc + G' \cdot \exp\left(\frac{\lambda}{2}\cos(2(\psi - \psi^*))\right), \tag{S42}$$

where the constant factor $\exp(\lambda/2)$ has been absorbed into the gain $G'$. For clarity, redefining $\lambda/2$ as $\lambda$ gives the $\pi$-periodic von Mises form

$$\Psi(\psi) = dc + G \cdot \exp\left(\lambda\cos(2(\psi - \psi^*))\right). \tag{S43}$$

***The manifold for roll-free head orientation is $\mathrm{T}^2$.*** As observed in Finkelstein et al. (2015), azimuth tuning of inverted bats is shifted by 180°, which makes the head direction tuning well modeled by a toroidal model. This suggests that the topological structure of "roll-free" head direction is $\mathrm{T}^2$. Here, we provide a mathematical proof supporting this finding.

For "roll-free" head orientation, the rotation matrix $R$ can be expressed as

$$R = R_z(\psi)R_y(\theta), \tag{S44}$$

where $\theta$ and $\psi$ represent pitch and azimuth angles, respectively. Using rotation matrices as defined in Eq. S4, $R$ expands to

$$R = \begin{bmatrix} \cos\psi\cos\theta & -\sin\psi & \cos\psi\sin\theta \\ \sin\psi\cos\theta & \cos\psi & \sin\psi\sin\theta \\ -\sin\theta & 0 & \cos\theta \end{bmatrix}. \tag{S45}$$

For simplification, we denote $a = \cos\theta$, $b = -\sin\theta$, $u = \cos\psi$, $v = -\sin\psi$, allowing $R$ be rewritten as

$$R = \begin{bmatrix} au & v & -bu \\ -av & u & bv \\ b & 0 & a \end{bmatrix}. \tag{S46}$$

Define $\mathcal{M} = \{R \in \mathrm{SO}(3) : r_{32} = 0\}$, where $r_{ij}$ $(i, j = 1, 2, 3)$ is the entry of $R$ on the $i^{\text{th}}$ row and $j^{\text{th}}$ column. Note that the $R$ matrix can be fully determined by the vectors $[a \;\; b]^\top \in \mathrm{S}^1$ and $[u \;\; v]^\top \in \mathrm{S}^1$. Define $f : \mathcal{M} \to \mathrm{T}^2$ via $f(R) = \{[a \;\; b]^\top, [u \;\; v]^\top\}$. To prove that $\mathcal{M}$ is diffeomorphic to $\mathrm{T}^2$ (i.e. $f$ is a diffeomorphism), we need to show that $f$ is bijective, $f$ and $f^{-1}$ are continuous, and $f$ is smooth.

Given a matrix $R \in \mathcal{M}$, we identify $a = r_{33}$, $b = r_{31}$, $u = r_{22}$ and $v = r_{12}$. Then the inverse map is

$$f^{-1}\left(\begin{bmatrix} a \\ b \end{bmatrix}, \begin{bmatrix} u \\ v \end{bmatrix}\right) = \begin{bmatrix} au & v & -bu \\ -av & u & bv \\ b & 0 & a \end{bmatrix} \in \mathcal{M}. \tag{S47}$$

Since $f$ has an explicit inverse, it is bijective.

Next, to prove $f$ is smooth and $f$ and $f^{-1}$ are continuous, we calculate the Jacobian of body velocity $\overline{\omega} = (R^{-1}\dot{R})^{\vee}$ as below:

$$\begin{aligned} \hat{\omega} &= R^{-1}\dot{R} \\ &= R^{-1}\left(\frac{\partial R}{\partial \theta}\dot{\theta} + \frac{\partial R}{\partial \psi}\dot{\psi}\right) \\ &= \begin{bmatrix} 0 & -\dot{\psi}\cos\theta & \dot{\theta} \\ \dot{\psi}\cos\theta & 0 & \dot{\psi}\sin\theta \\ -\dot{\theta} & -\dot{\psi}\sin\theta & 0 \end{bmatrix}, \end{aligned} \tag{S48}$$

which evaluates to

$$\overline{\omega} = \begin{bmatrix} -\dot{\psi}\sin\theta \\ \dot{\theta} \\ \dot{\psi}\cos\theta \end{bmatrix}. \tag{S49}$$

Since $\overline{\omega} = J^b(\theta,\psi)[\dot{\theta} \;\; \dot{\psi}]^\top$, the corresponding Jacobian matrix $J^b(\theta,\psi)$ is derived:

$$J^b(\theta,\psi) = \begin{bmatrix} 0 & -\sin\theta \\ 1 & 0 \\ 0 & \cos\theta \end{bmatrix}. \tag{S50}$$

$J^b(\theta,\psi)$ is full rank (rank = 2), indicating that there is no singularity and that $f$ is smooth with $f$ and $f^{-1}$ being continuous. Thus, we conclude that $f$ is a diffeomorphism, and "roll-free" rotations are globally parameterized by $\mathrm{T}^2$.

***Singularity exists in the gravity-based model.*** According to the parameterization in Angelaki et al. (2020), the coordinate of a point on the spherical manifold of 2D tilt is described by two angular parameters–$\alpha$ being the tilt angle and $\gamma$ being the tilt orientation. The remaining DOF is parameterized using $\theta$, the tilted azimuth that rotates the animal around the world-frame vertical axis. Therefore, the rotation matrix described by such parameterization can be calculated as

$$R = \exp(\hat{e_3}\theta)\exp(\hat{\omega}\alpha), \tag{S51}$$

where $\hat{\omega} = \begin{bmatrix}\sin\gamma & \cos\gamma & 0\end{bmatrix}^\top$. Term $\exp(\hat{e_3}\theta)$ denotes rotation around $e_3$ ($z$) axis by angle $\theta$, and $\exp(\hat{\omega}\alpha)$ denotes rotation around the axis defined by $\hat{\omega}$ by angle $\alpha$. Expanding Eq. S51, we get

$$R = \begin{bmatrix} r_{11} & r_{12} & r_{13} \\ r_{21} & r_{22} & r_{23} \\ r_{31} & r_{32} & r_{33} \end{bmatrix}, \tag{S52}$$

where

$$\begin{cases} r_{11} = \mathrm{c}\alpha\,\mathrm{c}\gamma\,\mathrm{c}(\gamma+\theta) + \mathrm{s}\gamma\,\mathrm{s}(\gamma+\theta) \\ r_{12} = (-1+\mathrm{c}\alpha)\,\mathrm{c}\gamma\,\mathrm{c}\theta\,\mathrm{s}\gamma - (\mathrm{c}^2\gamma + \mathrm{c}\alpha\,\mathrm{s}^2\gamma)\,\mathrm{s}\theta \\ r_{13} = \mathrm{s}\alpha\,\mathrm{c}(\gamma+\theta) \\ r_{21} = (-1+\mathrm{c}\alpha)\,\mathrm{c}\gamma\,\mathrm{c}\theta\,\mathrm{s}\gamma + (\mathrm{c}\alpha\,\mathrm{c}^2\gamma + \mathrm{s}^2\gamma)\,\mathrm{s}\theta \\ r_{22} = \mathrm{c}\theta\left(\mathrm{c}^2\gamma + \mathrm{c}\alpha\,\mathrm{s}^2\gamma\right) + (-1+\mathrm{c}\alpha)\,\mathrm{c}\gamma\,\mathrm{s}\gamma\,\mathrm{s}\theta \\ r_{23} = \mathrm{s}\alpha\,\mathrm{s}(\gamma+\theta) \\ r_{31} = -\mathrm{c}\gamma\,\mathrm{s}\alpha \\ r_{32} = -\mathrm{s}\alpha\,\mathrm{s}\gamma \\ r_{33} = \mathrm{c}\alpha. \end{cases} \tag{S53}$$

For simplicity, c denotes $\cos$ and s denotes $\sin$.

Define $\Psi : (\alpha,\gamma,\theta) \to \mathrm{SO}(3)$, where $\alpha \in [0,\pi]$, $\gamma \in [0,2\pi)$, $\theta \in [0,2\pi)$. If such parameterization using $(\alpha,\gamma,\theta)$ is a singularity-free parameterization of $\mathrm{SO}(3)$, $\Psi$ and $\Psi^{-1}$ should be smooth.

To test this, we can calculate the Jacobian matrix of $\Psi$ and see if it is singular. Following similar process in Eq. S48 and Eq. S49,

$$\overline{\omega} = \begin{bmatrix} -(\dot{\gamma}+\dot{\theta})\cos\gamma\sin\alpha - \dot{\alpha}\sin\gamma \\ \dot{\alpha}\cos\gamma - (\dot{\gamma}+\dot{\theta})\sin\alpha\sin\gamma \\ -\dot{\gamma} + (\dot{\gamma}+\dot{\theta})\cos\alpha \end{bmatrix}. \tag{S54}$$

and

$$J^b(\theta,\phi) = \begin{bmatrix} -\cos\gamma\sin\alpha & -\sin\gamma & -\cos\gamma\sin\alpha \\ -\sin\alpha\sin\gamma & \cos\gamma & -\sin\alpha\sin\gamma \\ \cos\alpha & 0 & -1+\cos\alpha \end{bmatrix}, \tag{S55}$$

whose determinant is $\sin\alpha$. This means that when $\alpha = 0$ or $\pi$, the Jacobian is singular, and $\Psi$ is not smooth any more. Therefore, the manifold of $\mathrm{SO}(3)$ is **not** $\mathrm{T}^2 \times \mathrm{S}^1$ or $\mathbb{R}^1 \times \mathrm{S}^1 \times \mathrm{S}^1$.

### Supplemental figures

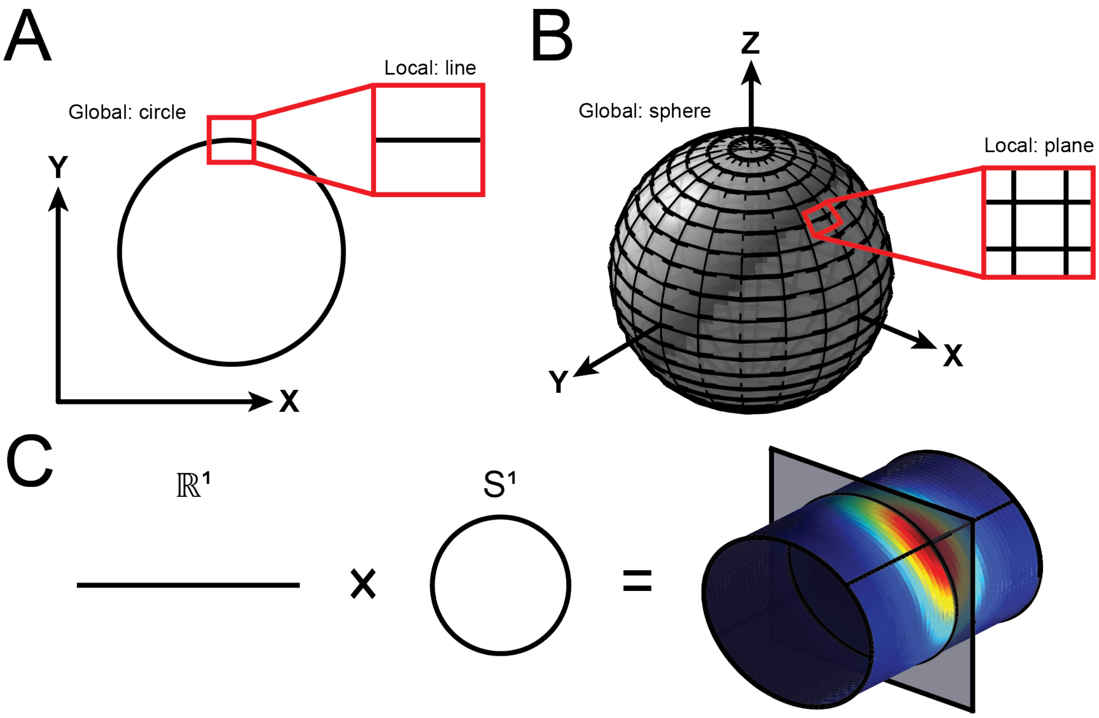


**Fig. S1. Geometric interpretation of manifolds. (A)** Intuitively speaking, lines and circles are 1D manifolds because they locally resemble lines and are parameterized by a single variable. **(B)** Likewise, planes and spheres are 2D manifolds because they locally resemble planes and are parameterized by two variables. **(C)** Manifolds describing distance and orientation of frontoparallel motion are $\mathbb{R}^1$ and $\mathrm{S}^1$, respectively. The joint configuration space, $\mathbb{R}^1 \times \mathrm{S}^1$, is geometrically the surface of an open-ended cylinder. Hypothetical neuronal responses illustrating joint tuning for distance and direction are shown as a heat map, with red (blue) hues indicate higher (lower) firing rates. Slices through the cylinder (transparent plane) are 1D circular manifolds describing motion direction and a strip along the long axis (black line) is a 1D linear manifold describing distance.

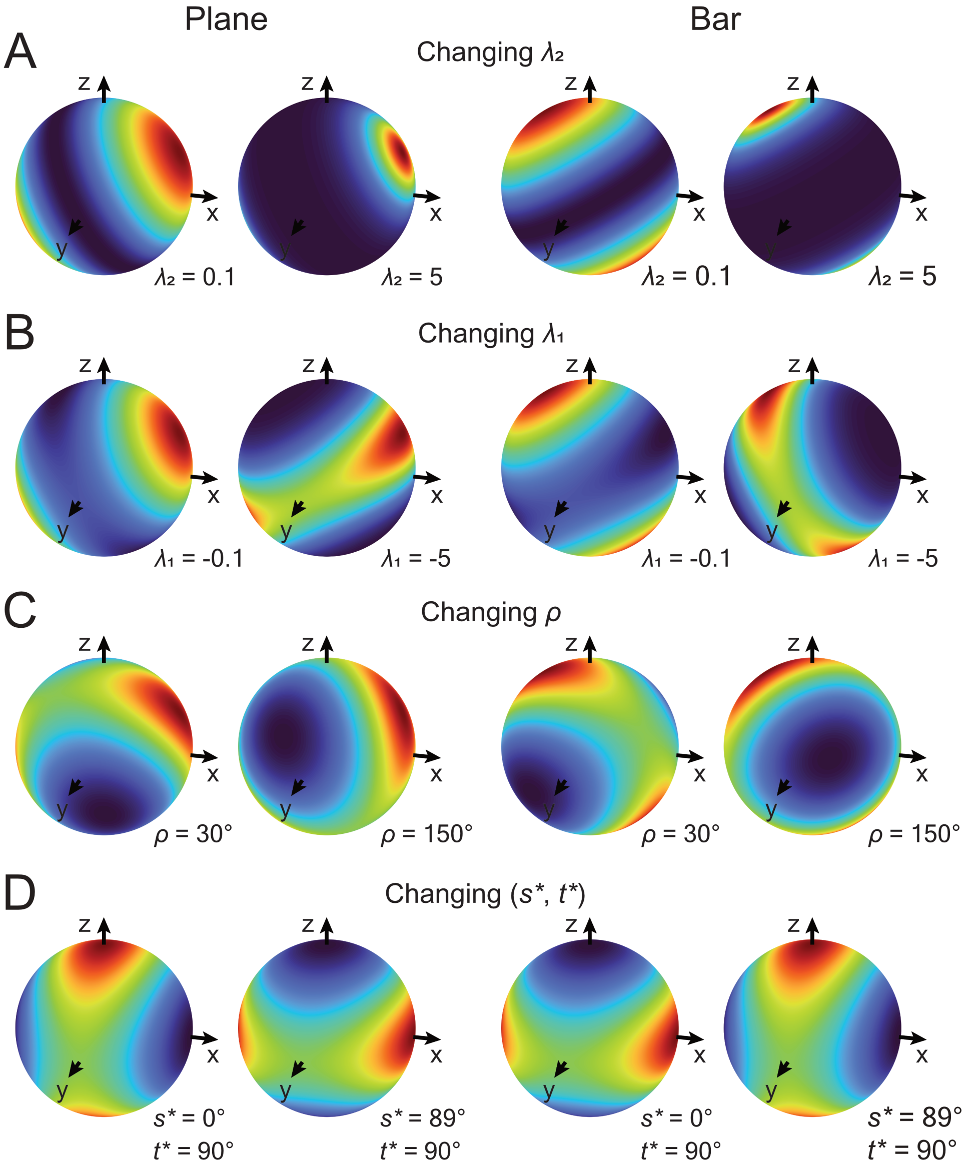


**Fig. S2. 3D orientation tuning curves corresponding to a plane and a bar when the Bingham function parameters are tuned to different values.** Left two columes: plane, where the central white dot denotes $s = 0^{\circ}$, and the white dashed line corresponds to $t = 0^{\circ}$; right two columes: bar, where $s = 0^{\circ}$ is along the bottom edge, and $t = 0^{\circ}$ along the left edge of the square. **(A)** Changing $\lambda_2$ ($\lambda_2 > 0$), which sets the bandwidth of Bingham function. Larger $\lambda_2$ leds to narrower tuning. Other parameters: $\lambda_1 = 0$, $\rho = 90^{\circ}$, $s^* = 30^{\circ}$, $t^* = 135^{\circ}$. **(B)** Changing $\lambda_1$ ($\lambda_1 < 0$), which determines the isotropy of Bingham function. More negative $\lambda_1$ leds to more anisotropy. Other parameters: $\lambda_2 = 0.7$, $\rho = 90^{\circ}$, $s^* = 30^{\circ}$, $t^* = 135^{\circ}$. **(C)** Changing $\rho$ ($0^{\circ} \leq \rho < 180^{\circ}$), which sets the rotation about the preferred slant-tilt axis. Other parameters: $\lambda_1 = -1.5$, $\lambda_2 = 0.7$, $s^* = 30^{\circ}$, $t^* = 135^{\circ}$. **(D)** Changing $s^*$ and $t^*$ ($0^{\circ} \leq s^* < 90^{\circ}$, $0^{\circ} \leq t^* < 360^{\circ}$), which define the preferred slant-tilt angles. Other parameters: $\lambda_1 = -1.5$, $\lambda_2 = 0.7$, $\rho = 90^{\circ}$.

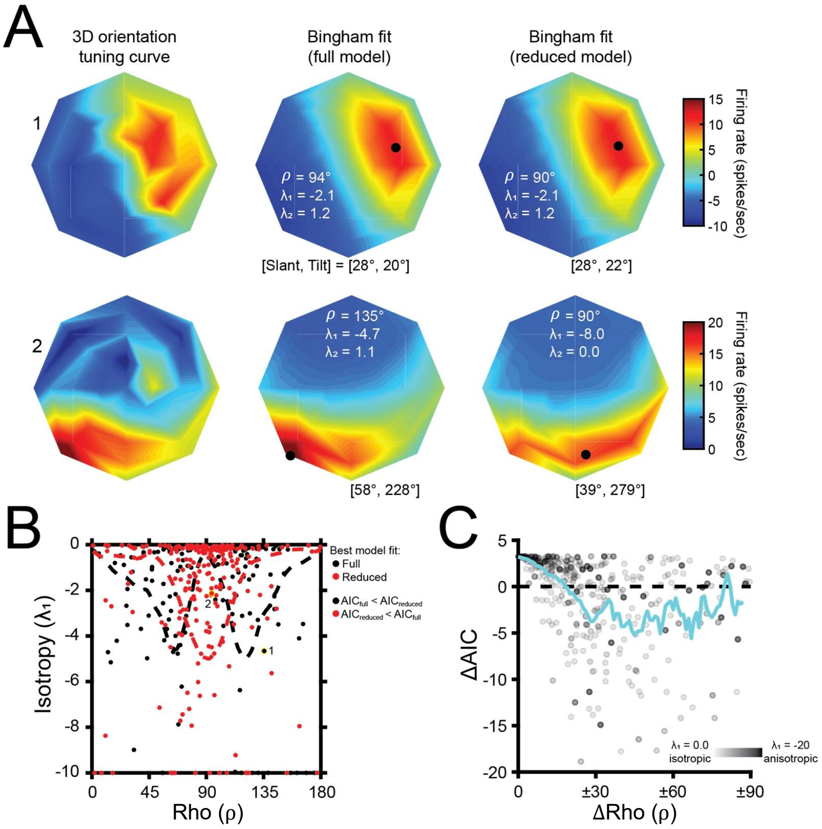


**Fig. S3. Comparing Bingham function fits with the full and reduced model. (A)** Slant–tilt tuning discs for two example CIP neurons. Slant is the radial axis, and tilt is the angular axis. Red hues indicate higher firing rates. Baseline-subtracted responses (left column) and Bingham fits using the full (center) and reduced (right) parameter set are shown. Black dots indicate the preferred slant–tilt. **(B)** Scatter plot comparing $\lambda_1$ (isotropy) vs $\rho$ from the full Bingham fits, for neurons better described by the full (black) or reduced (red) Bingham function. Dashed curve is a probability density function. **(C)** Scatter plot comparing the difference in AIC values for the full and reduced model vs the difference between $\rho$ from the full model and $90^\circ$. Positive $\Delta\mathrm{AIC}$ values indicate that the reduced model was a better fit of the data. Transparency is used it indicate the degree of isotropy ($\lambda_1$). A sliding window average (blue curve) is shown to illustrate trends.

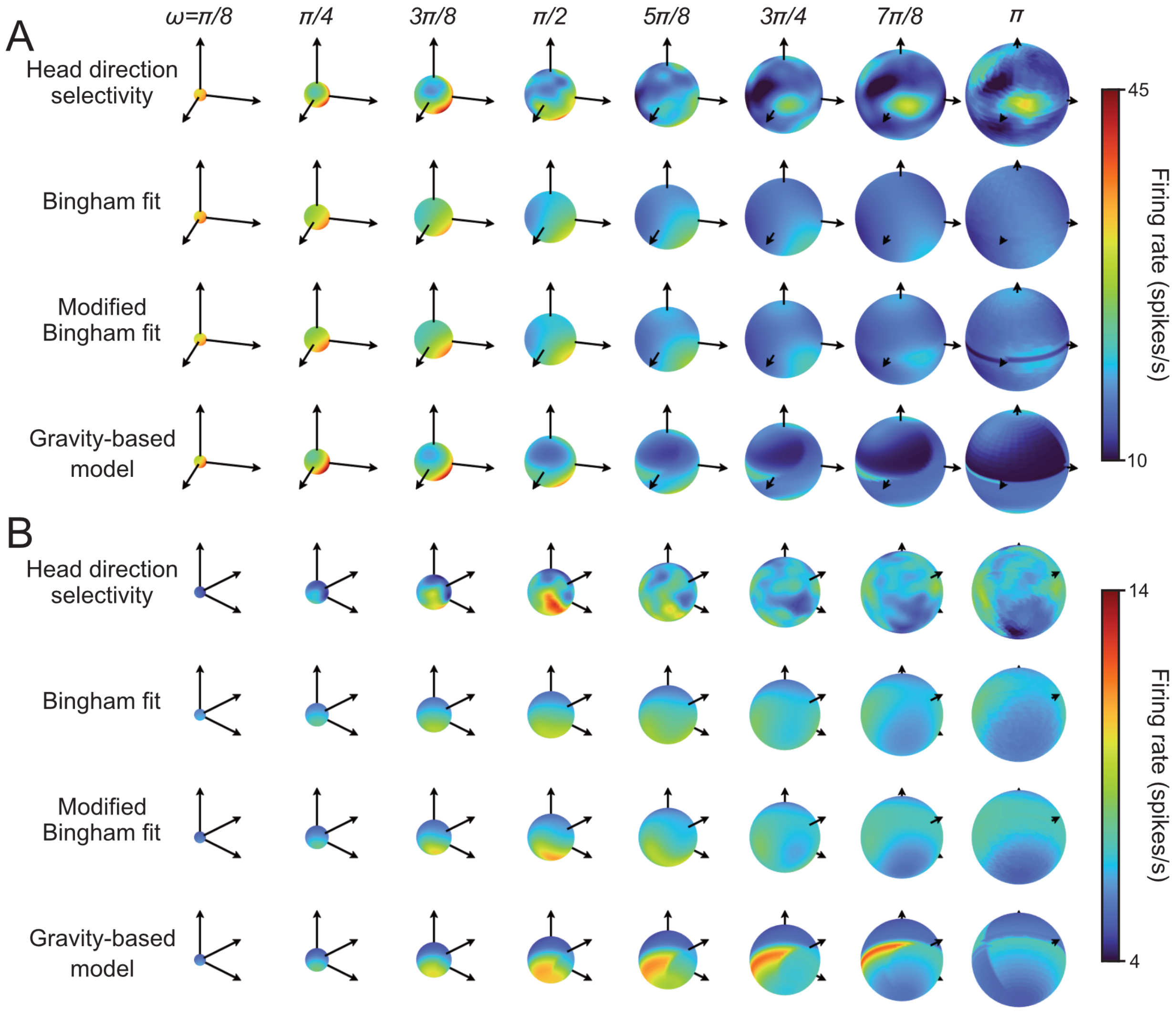


**Fig. S4. 3 DOF head direction tuning.** Responses of two neurons (**A, B**) recorded from the mouse cingulum fiber bundle for different head directions plotted using the shell visualization at $\omega = \pi/8$, $\pi/4$, $3\pi/8$, $\pi/2$, $5\pi/8$, $3\pi/4$, $7\pi/8$ and $\pi$ (row 1). Model fits using the Bingham function (row 2), the modified Bingham function (column 3) and gravity–based model (column 4) are also illustrated.

### Supplementary references

Alexa, M. Super-fibonacci spirals: Fast, low-discrepancy sampling of SO(3). In *2022 IEEE/CVF Conference on Computer Vision and Pattern Recognition (CVPR)*, pages 8281–8290, (2022). doi: 10.1109/CVPR52688.2022.00811.

Angelaki, D. E., Ng, J., Abrego, A. M., Cham, H. X., Asprodini, E. K., Dickman, J. D., and Laurens, J. (2020). A gravity-based three-dimensional compass in the mouse brain. *Nature communications*, 11(1):1855.

Baeg, E., Doudlah, R., Swader, R., Lee, H., Han, M., Kim, S.-G., Rosenberg, A., and Kim, B. (2021). Mri compatible, customizable, and 3D-printable microdrive for neuroscience research. *eNeuro*, 8(2). doi: 10.1523/ENEURO.0495-20.2021.

Burnham, K. P. and Anderson, D. R. *Model selection and multimodel inference : a practical information-theoretic approach*. Springer, New York, 2nd ed. edition, (2002). ISBN 0387953647.

Chang, T.-Y., Doudlah, R., Kim, B., Sunkara, A., Thompson, L. W., Lowe, M. E., and Rosenberg, A. (2020). Functional links between sensory representations, choice activity, and sensorimotor associations in parietal cortex. *Elife*, 9:e57968.

Doudlah, R., Chang, T.-Y., Thompson, L. W., Kim, B., Sunkara, A., and Rosenberg, A. (2022). Parallel processing, hierarchical transformations, and sensorimotor associations along the 'where'pathway. *Elife*, 11:e78712.

Finkelstein, A., Derdikman, D., Rubin, A., Foerster, J. N., Las, L., and Ulanovsky, N. (2015). Three-dimensional head-direction coding in the bat brain. *Nature (London)*, 517(7533):159–164.

Jones, E. *The Thalamus*. Number v. 1 in The Thalamus. Cambridge University Press, (2007). ISBN 9780521858816. URL https://books.google.com/books?id=EtTvsgEACAAJ.

Kim, B., Kenchappa, S. C., Sunkara, A., Chang, T.-Y., Thompson, L., Doudlah, R., and Rosenberg, A. (2019). Real-time experimental control using network-based parallel processing. *eLife*, 8.

Koditschek, D. E. Application of a new Lyapunov function to global adaptive attitude tracking. In *Proceedings of the IEEE Conference on Decision and Control*, pages 63–68, (1988).

Laurens, J., Kim, B., Dickman, J. D., and Angelaki, D. E. (2016). Gravity orientation tuning in macaque anterior thalamus. *Nature neuroscience*, 19(12):1566–1568.

Rosenberg, A., Cowan, N. J., and Angelaki, D. E. (2013). The visual representation of 3D object orientation in parietal cortex. *The Journal of neuroscience*, 33(49):19352–19361.

Stevens, K. A. (1983). Slant-tilt: The visual encoding of surface orientation. *Biol Cybern*, 46(3):183–195.
